# Thalamic integration of basal ganglia and cerebellar circuits during motor learning

**DOI:** 10.1101/2024.10.31.621388

**Authors:** Richard H. Roth, Michael A. Muniak, Charles J. Huang, Fuu-Jiun Hwang, Yue Sun, Cierra Min, Tianyi Mao, Jun B. Ding

**Affiliations:** Department of Neurosurgery, Stanford University, Stanford, CA 94305, USA; Aligning Science Across Parkinson’s (ASAP) Collaborative Research Network, Chevy Chase, MD 20815, USA; Vollum Institute, Oregon Health and Science University, Portland, OR 97239, USA; Department of Neurology and Neurological Sciences, Stanford University, Stanford, CA 94305, USA; The Phil & Penny Knight Initiative for Brain Resilience at the Wu Tsai Neurosciences Institute, Stanford University, Stanford, CA 94305, USA

**Keywords:** motor learning, thalamus, basal ganglia, cerebellum, circuit plasticity, circuit convergence, cell mapping, multi-site photometry

## Abstract

The ability to control movement and learn new motor skills is one of the fundamental functions of the brain. The basal ganglia (BG) and the cerebellum (CB) are two key brain regions involved in controlling movement, and neuronal plasticity within these two regions is crucial for acquiring new motor skills. However, how these regions interact to produce a cohesive unified motor output remains elusive. Here, we discovered that a subset of neurons in the motor thalamus receive converging synaptic inputs from both BG and CB. By performing multi-site fiber photometry in mice learning motor tasks, we found that motor thalamus neurons integrate BG and CB signals and show distinct movement-related activity. Lastly, we found a critical role of these thalamic neurons and their BG and CB inputs in motor learning and control. These results identify the thalamic convergence of BG and CB and its crucial role in integrating movement signals.

**Highlights:**

- Individual neurons in motor thalamus receive converging synaptic input from SNr and DCN projections.
- Thalamic neurons with SNr and DCN input are concentrated at the border between VM and VAL thalamic nuclei.
- Thalamic neurons functionally integrate SNr and DCN activity and adapt with motor learning.
- Thalamic neurons and their inputs from SNr and DCN are critical for learning and executing motor tasks.

## Introduction

The learning and execution of motor skills require coordinated activity across multiple regions in the brain. Among these, the basal ganglia (BG) and cerebellar (CB) systems play essential roles in motor control and learning. The BG is involved in voluntary movement, action selection, and habit formation^1,2^, while the CB performs sensory integration, motor timing, and motor predictions^3–5^. Despite the importance of these two major brain circuits’ individual roles in motor function, how they interact with each other to produce cohesive motor outputs remains elusive^6^. Particularly, where in the brain the BG and CB regions first interact and how critical this interplay is for motor control remains a critical question.

Anatomically, the thalamus represents a common target for BG output from the Substantia Nigra pars reticulata (SNr) and CB output from the deep cerebellar nuclei (DCN) and in turn sends projections to the cortex^7–10^. The motor thalamus is conventionally divided into a distinct BG recipient zone and CB recipient zone, as BG projections target the ventral medial thalamus (VM) and the anterior part of the ventral anterior-lateral complex (VAL, or VA), whereas CB projections target the posterior part of VAL (or VL)^11–17^. This convention views the thalamus as a passive relay station for BG and CB inputs, suggesting that any functional interaction of these motor circuits occurs at the cortical level^18,19^.

The notion that BG and CB circuits only interact at the cortical level, however, has been challenged by the discovery of subcortical connections between BG and CB^20,21^. Emerging anatomical and physiological evidence further suggests that, contrary to this conventional view, the thalamus does receive overlapping inputs from BG and DCN and may functionally integrate these diverse motor signals^18,22– 24^. Specifically, regions within VM and VAL appear to receive projections from both BG and CB^25–28^. Currently, the exact location and extent of thalamic neurons receiving both BG and CB inputs and their role in functionally integrating these signals during motor control and learning remain unknown.

In this study, we identified and labeled a substantial population of neurons in the thalamus that receive converging synaptic inputs from the SNr and DCN using transsynaptic viral tracing and slice electrophysiology approaches. Mapping these neurons to the Allen Brain Atlas^29^ and Franklin-Paxinos Atlas^30,31^, we found that while distributed across multiple thalamic nuclei, the highest density of dual-input neurons is located at the border between VM and VAL. By training animals on multiple motor tasks and simultaneously recording neuronal activity in SNr, DCN, and dual-input thalamic neurons, we found that thalamic neurons functionally integrate SNr and DCN activity and differentially adapt their activity throughout motor learning. Lastly, inhibiting SNr or DCN projections in the thalamus or selectively lesioning thalamic neurons receiving both inputs impaired motor performance and learning. Together, our results suggest that the thalamus is a key node for integrating BG and CB motor signals and that their anatomical convergence onto individual neurons in the thalamus is critical for learning and executing motor skills.

## Results

### Thalamus neurons receive converging inputs from SNr and DCN projections

The thalamus is a major target of basal ganglia outputs from the SNr and cerebellar outputs from the DCN. To measure the convergence of these projections, we first used a two-color anterograde viral tracing approach to label projection neurons and their axons from the SNr and the DCN. Specifically, we injected one adeno-associated virus (AAV) expressing a red fluorescent protein (AAV-tdTomato) in the SNr and another AAV expressing GFP (AAV-GFP) in the DCN (**Fig. 1A-B**). We aimed for a comprehensive labeling of SNr and DCN neurons since a majority of neuronal subpopulations in these regions send projections to motor nuclei of the thalamus (VM/VAL)^4,32–38^. In these mice, we replicated previous findings demonstrating that SNr projections are present in VM and the anterior part of VAL (BG recipient zone), while DCN axons are located in the posterior part of VAL (CB recipient zone). However, we also observed a substantial overlap of SNr and DCN axons in the thalamus, especially at the VM and VAL border (**Fig. 1C**).

**Figure 1:**
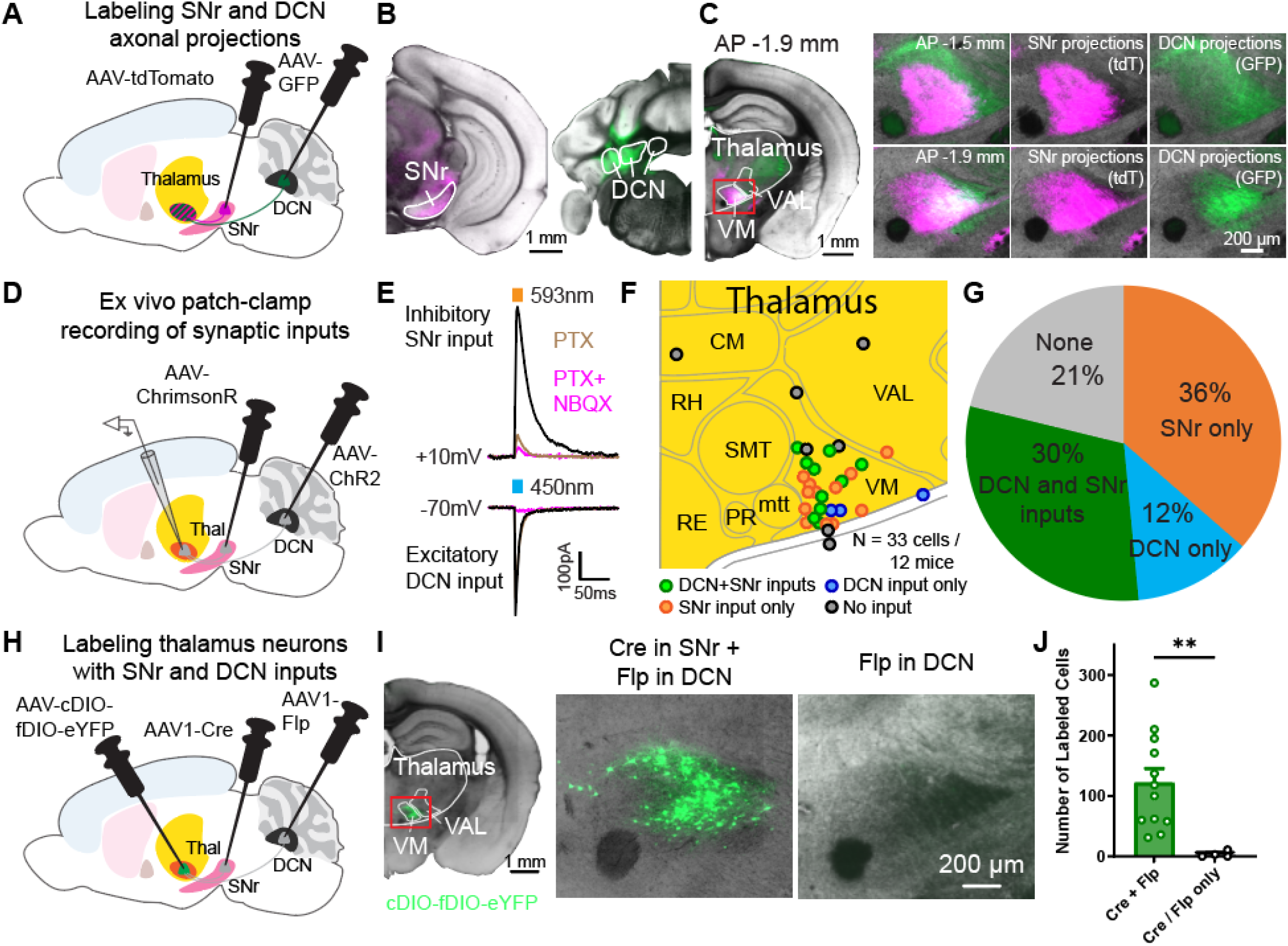
Neurons in motor thalamus receive converging synaptic inputs from SNr and DCN projections. (A) Schematic diagram of virus injections for viral tracing of SNr and DCN axonal projections. (B) Coronal sections showing AAV-tdTomato (magenta) expression in SNr (left) and AAV-GFP (green) in DCN (right). Scale bar, 1 mm. (C) Coronal sections showing axonal projections from ipsilateral SNr and contralateral DCN in the thalamus. Left: Hemisphere with SNr axons expressing AAV-tdTomato (magenta) and DCN axons expressing AAV-GFP (green) at 1.9 mm posterior of Bregma. Scale bar, 1 mm. Right: Enlarged images of motor thalamus (red rectangle in image on the left) at 1.5 mm (top) and 1.9 mm (bottom) posterior of Bregma. Scale bar, 200 μm. (D) Schematic diagram of virus injections for dual-color optogenetic stimulation and whole-cell patch clamp experiments. (E) Example whole-cell patch clamp recording traces of oIPSC and oEPSC responses in a thalamic neuron to stimulation of SNr inputs with ChrimsonR and DCN inputs with ChR2. oIPSCs (black) were blocked by PTX application (dark magenta), oEPSCs (black) were blocked by NBQX application (light magenta). (F) Location of recorded neurons, colored by their input type. Recorded neurons from different AP sections (1.3-1.9 mm posterior of Bregma) shown on one map for clarity. (G) Fraction of neurons receiving functional SNr and/or DCN inputs. Data in F and G are from a mix of mice expressing ChrimsonR in SNr and ChR2 in DCN or vice versa. N=33 cells from 12 mice. (H) Schematic diagram of virus injections for transsynaptic viral tracing of SNr and DCN neurons projecting onto thalamic neurons. (I) Coronal section showing neurons in the thalamus receiving dual inputs from SNr and DCN. Left: Hemisphere with neurons expressing AAV-cDIO-fDIO-eYFP (green) in a mouse injected with AAV1-Cre in the SNr and AAV1-Flp in the DCN. Scale bar, 1 mm. Middle: Enlarged image of motor thalamus (red rectangle in image on the left). Right: Motor thalamus in control mice injected with only AAV1-Cre or AAV1-Flp in SNr/DCN and AAV-cDIO-fDIO-eYFP in thalamus. (J) Quantification of labeled cells from two coronal sections in mice expressing AAV-cDIO-fDIO-eYFP in thalamus and AAV1-Cre in SNr and AAV1-Flp in DCN (or vice versa) and control mice. Bars denote mean and circles individual mice. Error bars, SEM. Cre+Flp: 122.1±23.13 cells, n=12 mice; Cre/Flp only: 3.5±2.6 cells, n=4 mice. p=0.001, Mann-Whitney test. ^∗∗^p<0.01

To probe whether these overlapping projections lead to individual thalamic neurons receiving functional synaptic inputs from both the SNr and DCN, we performed whole-cell patch-clamp experiments in the thalamus while optogenetically stimulating SNr or DCN inputs. Specifically, we expressed blue light-activatable channelrhodopsin (AAV-ChR2) in the DCN and an orange light-activatable opsin (AAV-ChrimsonR) in the SNr, or vice versa (**Fig. 1D, S1A-B**). This optogenetic approach allowed us to independently activate excitatory DCN projections using 450nm blue light or inhibitory SNr projections with 593nm orange light in acute brain slices. Within individual thalamic neurons, we recorded outward currents when stimulating SNr projections which were blocked by the GABA_A_ receptor blocker Picrotoxin (PTX), and inward currents upon stimulation of DCN projections, which were blocked by the AMPA receptor antagonist NBQX (**Fig. 1E**). Out of the 33 neurons recorded within the ventral thalamus (**Fig. 1F**), ∼30% of neurons received dual inputs from DCN and SNr, ∼36% received only SNr input, and ∼12% received only DCN input (**Fig. 1G, S1C-D**). The majority of neurons with dual inputs appeared to be located within the VM (**Fig. 1F**).

To further identify thalamic neurons receiving inputs from both SNr and DCN projections, we used an orthogonal approach that combined an anterograde transsynaptic viral labeling strategy using AAV1^39^ with an intersectional recombinase system using Cre and Flp^40^. By injecting AAV1-Cre in the SNr, AAV1-Flp in the DCN, and a Cre- and Flp-dependent fluorescent reporter (AAV-cDIO-fDIO-eYFP) in the thalamus (**Fig. 1H**), we could specifically label thalamic neurons with SNr and DCN inputs (**Fig. 1I**). In control mice expressing the reporter virus in motor thalamus with either only AAV1-Cre or only AAV1-Flp presynaptically, no labeled neurons were found (**Fig. 1I-J**). Together, these findings demonstrate that individual neurons in the thalamus receive functional converging synaptic inputs from both DCN and SNr.

### Most thalamic neurons with SNr and DCN inputs are located in VM and VAL

Next, we used this Cre- and Flp-dependent viral strategy to map the precise location of neurons with dual SNr and DCN inputs unbiasedly. We, therefore, employed algorithms to compare these cells across animals and map their location to the Allen Brain Atlas^29^ and Franklin-Paxinos Atlas^7,30,31^. Specifically, we sectioned and imaged the whole brain of mice with eYFP expression in dual-input thalamic neurons and aligned the sections for 3D reconstruction (**Fig. 2A**). Reconstructed brains were then registered to the Allen Brain Atlas CCFv3 using affine transformation based on stereotypic fiducials (see methods, **Fig. 2B**). We used this alignment and registration to map the location of labeled neurons and quantify which brain regions and, more specifically, which thalamic nuclei contained dual-input neurons (**Fig. 2C**). The same procedure was applied for registering brain sections to the Franklin-Paxinos Atlas (**Fig. S2**). Across nine brains with the AAV-cDIO-fDIO-eYFP reporter injected at three injection sites covering the ventral thalamus, along with AAV1-Cre in the SNr and AAV1-Flp in the DCN (or vice versa), we labeled an average of 1353 ± 264 cells and found that the majority of dual-input neurons were located within the thalamus (**Fig. 2D-F, S2A-C**). A substantial number of labeled neurons was also found in the Zona Incerta. These data indicate that the motor thalamus is a key area of convergence for BG and CB inputs and, therefore, likely plays a crucial role in functionally integrating signals from both circuits during motor control and motor learning.

**Figure 2:**
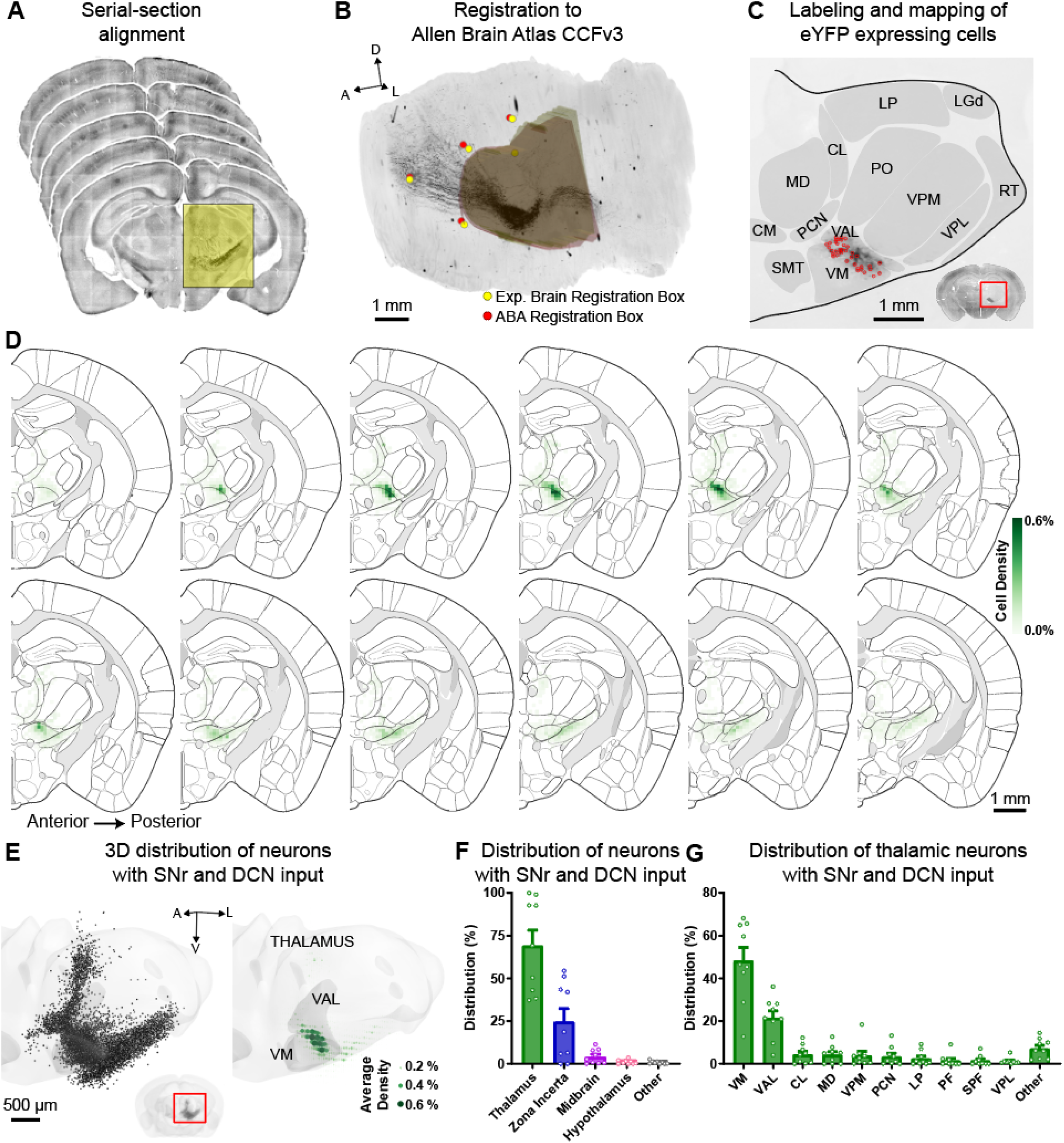
Most thalamic neurons with SNr and DCN inputs are located in VM and VAL. (A) Example serial brain sections from mice expressing AAV1-Cre or AAV1-Flp in SNr/DCN and AAV-cDIO-fDIO-eYFP in thalamus that are aligned for 3D reconstruction following imaging. (B) Aligned sections are registered to the Allen Brain Atlas CCFv3 using a sequence of affine transformations derived from matching brain landmark fiducials and coronal bounding boxes around the thalamus. Yellow circles/3D volume show a set of these registration landmarks for the experimental brain and red circles/3D volume show the corresponding and aligned landmarks from the Allen Brain Atlas (ABA). (C) Example section showing labeled neurons (red circles) mapped to thalamic nuclei from the Allen Brain Atlas. Inset shows the entire coronal section with enlarged region marked with a red rectangle. (D) Serial 2D coronal sections at 100-μm intervals of the Allen Brain Atlas. The distribution of labeled neurons with SNr and DCN inputs is shown as the average relative density in 100 × 100 μm bins (green, see methods) from n=9 mice. (E) 3D projection of the thalamus with individual labeled neurons from all 9 mice overlayed (left) and the average relative density of labeled neurons in 100 × 100 × 100 μm bins (right). Thalamus volume in light gray, VM and VAL highlighted in dark gray. Green circles denote average relative density. Inset shows 3D reconstruction of the entire brain with the enlarged region marked with a red rectangle. (F) Distribution of neurons with SNr and DCN inputs across brain regions. Bars denote mean and circles individual mice. Error bars, SEM. n=12,181 cells from 9 mice. (G) Distribution of neurons with SNr and DCN inputs within the thalamus. Individual thalamic nuclei with an average of ≥1% of labeled cells are listed, remaining nuclei are combined as “other”. Bars denote mean and circles individual mice. Error bars, SEM. n=7,514 cells from 9 mice.

Within the thalamus, the motor nuclei VM and VAL (more specifically the VL when using the Franklin-Paxinos Atlas) harbored most of the neurons with SNr and DCN input; though, intralaminar thalamic nuclei, including the central lateral (CL), paracentral (PCN), and parafascicular (PF) nuclei, also contained a small proportion of these dual-input neurons (**Fig. 2G, S2D-F**). We observed the highest density of dual-input thalamic neurons at the VM and VAL/VL border (**Fig. 2D, S2E-F**). These findings are consistent with the idea that BG and CB signal integration in the thalamus occurs at border regions between the BG and CB recipient areas^22^, which corresponds to the areas with highest overlap of axonal projections from SNr and DCN (**Fig. 1C**).

### Thalamic neurons functionally integrate SNr and DCN activity

Neurons in the SNr, DCN, and the motor thalamus play an important functional role in movement control and motor learning^1,4,41–47^. We therefore investigated whether thalamic neurons receiving converging synaptic inputs from the SNr and DCN also functionally integrate these signals and contribute to motor learning. Using multi-site Ca^2+^ fiber photometry, we simultaneously recorded the activity of SNr and DCN as well as thalamic neurons with dual SNr and DCN inputs, while training mice on head-fixed lever-pushing and head-fixed locomotion tasks (**Fig. 3A, S3A**). To do so, we expressed soma-targeted GCaMP8m (AAV-Ribo-1L-GCaMP8m) in the SNr and DCN and combined this with the transsynaptic AAV1 labeling approach to express GCaMP8m specifically in dual-input thalamic neurons (**Fig. 3A-B**). To avoid influence of Ca^2+^ signals from SNr and DCN axonal projections into thalamus recordings, we used soma-targeted GCaMP8m in the SNr and DCN^48^.

**Figure 3:**
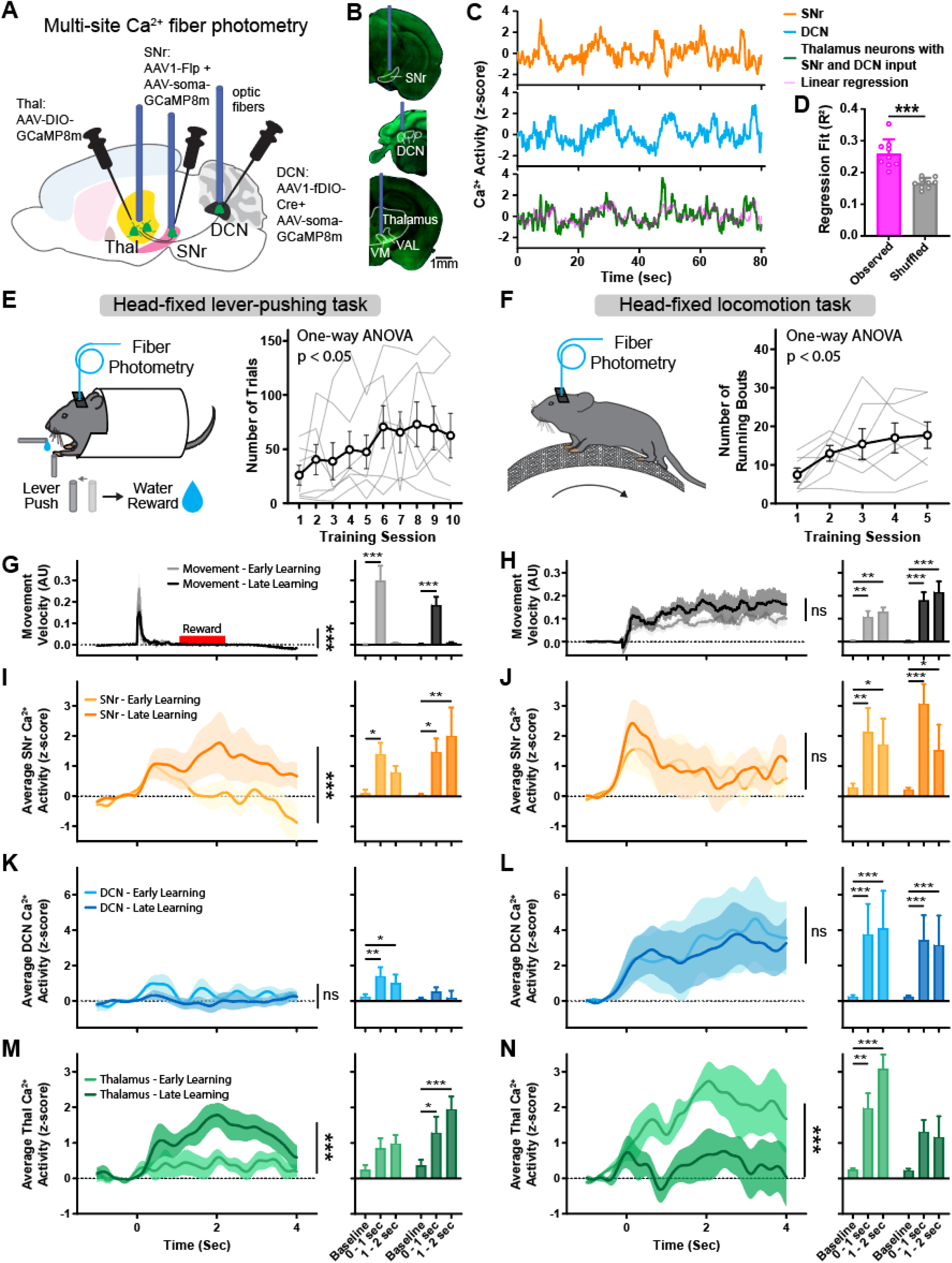
Functional integration of BG and CB activity in the thalamus during motor learning. (A) Schematic diagram of virus injections and optic fiber placement for multi-site fiber photometry of Ca^2+^ activity in SNr, DCN, and thalamus neurons with dual inputs. (B) Example coronal sections showing AAV-soma-GCaMP8m (SNr and DCN, green) and AAV-GCaMP8m (thalamus, green) expression and optic fiber placement. Scale bar, 1 mm (C) Example fiber photometry recording traces from SNr (orange), DCN (blue), and dual-input thalamus neurons (green) in one mouse. Predicted thalamus signal based on linear regression of SNr and DCN signals (A × Ca^2+^ _SNr_ + B × Ca^2+^_DCN_ + C = Ca^2+^ _Thalamus_) shown in magenta. (D) Regression fit (R^2^) of predicted thalamus signal calculated from linear regression of random 135 sec time intervals (see methods) from observed data (magenta) and predicted signal calculated from control shuffled data with SNr and DCN signals randomly shifted in time by up to ±67.5 sec relative to the thalamus signal (grey). Bars denote mean and circles individual mice. Error bars, SEM. R^2^ of observed data: 0.26±0.015; R^2^ of shuffled data 0.17±0.005, n=9 mice. p<0.001, paired T-test. (E) Left: Schematic of mouse performing head-fixed lever-pushing task. Right: Behavior performance of mice learning the lever-pushing task over the course of 10 days. Thin grey lines represent individual mice and bold line denotes mean. Error bars, SEM. n=7 mice. p=0.04, repeated measure one-way ANOVA. (F) Left: Schematic of mouse performing head-fixed locomotion task. Right: Behavior performance of mice learning the locomotion task over the course of 5 days. Thin grey lines represent individual mice and bold line denotes mean. Error bars, SEM. n=7 mice. p=0.02, repeated measure one-way ANOVA. (G) Left: Average velocity of lever movements aligned to movement onset during early (grey) and late (black) training sessions. Time window of water reward delivery shown in red. Solid lines represent mean and shaded areas denote SEM. n=7 mice. p<0.001, two-way ANOVA interaction. Right: Average maximal movement velocity at baseline, during movement (0-1 sec), and after movement (1-2 sec). Bars denote mean. Error bars, SEM. Two-way ANOVA with Holm-Sidak multiple comparisons test. (H) Left: Average velocity of wheel movements aligned to movement onset during early (grey) and late (black) training sessions. Solid lines represent mean and shaded areas denote SEM. n=7 mice. p=0.883, two-way ANOVA interaction. Right: Average maximal movement velocity at baseline, at movement onset (0-1 sec), and during continuous movement (1-2 sec). Bars denote mean. Error bars, SEM. Two-way ANOVA with Holm-Sidak multiple comparisons test. (I) Left: Average z-scored Ca^2+^ activity in SNr aligned to lever movement onset during early (light orange) and late (dark orange) training sessions. Solid lines represent mean and shaded areas denote SEM. n=7 mice. p<0.001, two-way ANOVA interaction. Right: Average maximal z-scored Ca^2+^ activity at baseline, during movement (0-1 sec), and after movement (1-2 sec). Bars denote mean. Error bars, SEM. Two-way ANOVA with Holm-Sidak multiple comparisons test. (J) Left: Average z-scored Ca^2+^ activity in SNr aligned to wheel movement onset during early (light orange) and late (dark orange) training sessions. Solid lines represent mean and shaded areas denote SEM. n=7 mice. p=0.709, two-way ANOVA interaction. Right: Average maximal z-scored Ca^2+^ activity at baseline, at movement onset (0-1 sec), and during continuous movement (1-2 sec). Bars denote mean. Error bars, SEM. Two-way ANOVA with Holm-Sidak multiple comparisons test. (K) Left: Average z-scored Ca^2+^ activity in DCN aligned to lever movement onset during early (light blue) and late (dark blue) training sessions. Solid lines represent mean and shaded areas denote SEM. n=7 mice. p=0.898, two-way ANOVA interaction. Right: Average maximal z-scored Ca^2+^ activity at baseline, during movement (0-1 sec), and after movement (1-2 sec). Bars denote mean. Error bars, SEM. Two-way ANOVA with Holm-Sidak multiple comparisons test. (L) Left: Average z-scored Ca^2+^ activity in DCN aligned to wheel movement onset during early (light blue) and late (dark blue) training sessions. Solid lines represent mean and shaded areas denote SEM. n=7 mice. p=0.976, two-way ANOVA interaction. Right: Average maximal z-scored Ca^2+^ activity at baseline, at movement onset (0-1 sec), and during continuous movement (1-2 sec). Bars denote mean. Error bars, SEM. Two-way ANOVA with Holm-Sidak multiple comparisons test. (M) Left: Average z-scored Ca^2+^ activity in thalamus neurons with SNr and DCN inputs aligned to lever movement onset during early (light green) and late (dark green) training sessions. Solid lines represent mean and shaded areas denote SEM. n=7 mice. p<0.001, two-way ANOVA interaction. Right: Average maximal z-scored Ca^2+^ activity at baseline, during movement (0-1 sec), and after movement (1-2 sec). Bars denote mean. Error bars, SEM. Two-way ANOVA with Holm-Sidak multiple comparisons test. (N) Left: Average z-scored Ca^2+^ activity in thalamus neurons with SNr and DCN inputs aligned to wheel movement onset during early (light green) and late (dark green) training sessions. Solid lines represent mean and shaded areas denote SEM. n=7 mice. p<0.001, two-way ANOVA interaction. Right: Average maximal z-scored Ca^2+^ activity at baseline, at movement onset (0-1 sec), and during continuous movement (1-2 sec). Bars denote mean. Error bars, SEM. Two-way ANOVA with Holm-Sidak multiple comparisons test. ^∗^p<0.05, ^∗∗^p<0.01, ^∗∗∗^p<0.001

Overall, the population activity of thalamic neurons with inputs from SNr and DCN showed correlation with the activity in these input regions (**Fig. 3C**). To test the relationship between neuronal activity in the thalamus and its input regions, we performed a multiple linear regression analysis to predict thalamus activity signal based on SNr and DCN activity (**Fig. 3C**, magenta). Overall, this predicted thalamus signal corresponded closely with the actual thalamus signal. We then quantified the specificity of this property by creating a shuffled control condition, in which SNr and DCN signals were randomly shifted in time relative to the thalamus signal. As a result, we noted a significant drop in the quality and fit of the linear regression (**Fig. 3D**).

### Thalamic activity adapts with motor learning

We trained mice on two commonly used motor tasks to test how SNr, DCN, and dual-input thalamus neuron activity adapts throughout motor learning. We selected two distinct behaviors – a head-fixed lever-pushing task (**Fig. 3E**) and a head-fixed locomotion task (**Fig. 3F**) – that differ in their motor demands and engage cortico-basal ganglia and cerebellar circuits^49–53^. In the lever-pushing task, water-restricted mice pushed a joystick lever across a set distance threshold to receive a water reward in a self-initiated manner. Mice learned to perform this task by increasing the number of lever-pushes over the course of ten 15-minute training sessions (**Fig. 3E, S3B-C**). Comparing the early learning phase (first three training sessions) with the late phase (last three training sessions), we found a significant decrease in lever movement velocity (**Fig. 3G**).

When examining the neuronal activity, we found that the lever-pushing task predominantly activated the SNr, with activity aligned to movement onset and during reward delivery (**Fig. 3I**). SNr activity increased significantly throughout motor learning, particularly during reward delivery. DCN also showed activity linked to movement and reward but only during the early training sessions (**Fig. 3K**). The activity of thalamic neurons receiving inputs from both the SNr and DCN increased significantly over the course of learning the lever-pushing task, with movement and reward periods in the late training sessions showing significantly higher activity (**Fig. 3M**).

Comparing neuronal activity between successful, rewarded lever pushes and failed, unrewarded lever-push trials where the mice did not push the lever across the threshold, we found that SNr and thalamic activity was markedly stronger in rewarded trials (**Fig S3F-K**). Following motor learning, SNr and thalamic activity increased in both rewarded and unrewarded trials.

The head-fixed locomotion task involved mice voluntarily running on a cylindrical treadmill, and performance was quantified in the number of running bouts throughout five 15-minute training sessions (**Fig. 3F**). We observed no obvious differences in running velocity between early (first two) training sessions and late (last two) training sessions (**Fig. 3H**). During running bouts, activity in SNr peaked at motion onset and was sustained during ongoing locomotion (**Fig. 3J**). In contrast to the lever-pushing task, this task also involved strong activation of neurons in the DCN throughout the movement period (**Fig. 3L**). Notably, neither SNr nor DCN activity adapted over the course of this locomotion task. In contrast, the activity of thalamic neurons with SNr and DCN inputs was higher than baseline during early training sessions and decreased significantly throughout learning (**Fig. 3N**).

These distinct adaptations of SNr, DCN, and dual-input thalamic neurons across two different motor tasks suggest that thalamic neurons do not simply follow the activity of their input regions SNr and DCN but instead perform an active role in integrating and relaying these signals.

### Silencing SNr and DCN inputs and convergence impairs motor behaviors

The distinct adaptations observed in converging SNr-thalamus and DCN-thalamus circuits suggest that these projections and their integration by dual-input motor thalamic neurons may play a critical role in movement control and motor learning. To test whether SNr and DCN inputs to the thalamus are required for learning and executing motor tasks, we first trained mice on the head-fixed lever-pushing task and optogenetically silenced either SNr or DCN projections to the thalamus in well-trained mice (**Fig. 4A, S4A-B**). We expressed the opsin eOPN3 in the SNr of VGAT-cre mice or in the DCN of mice injected with retrograde-AAV-Cre in the thalamus. We then implanted an optic fiber in their target region within the thalamus to specifically block synaptic transmission of SNr or DCN projections in the thalamus (**Fig. 4B-I**)^54^. During two optogenetic silencing sessions following head-fixed lever-pushing training, mice were first allowed to perform the task during a 2-minute baseline period without inhibition. SNr or DCN projections were then inhibited for 3 minutes, followed by a post-inhibition recovery period. When inhibiting SNr inputs to the thalamus, we observed a significant drop in the number of successful pushes during stimulation, which recovered to baseline levels within 1 min following inhibition (**Fig. 4D and S4C**). Control mice injected with mCherry did not show any changes. Similarly, the total number of pushes did not change with inhibition (**Fig. S4D**), suggesting that the learned behavior was specifically impaired. When inhibiting DCN inputs to the thalamus, we did not observe any change in performance on the lever-pushing task (**Fig. 4H, S4E-F**), consistent with our Ca^2+^ photometry data showing that executing this task predominantly involves SNr activity (**Fig. 3I-K**).

**Figure 4:**
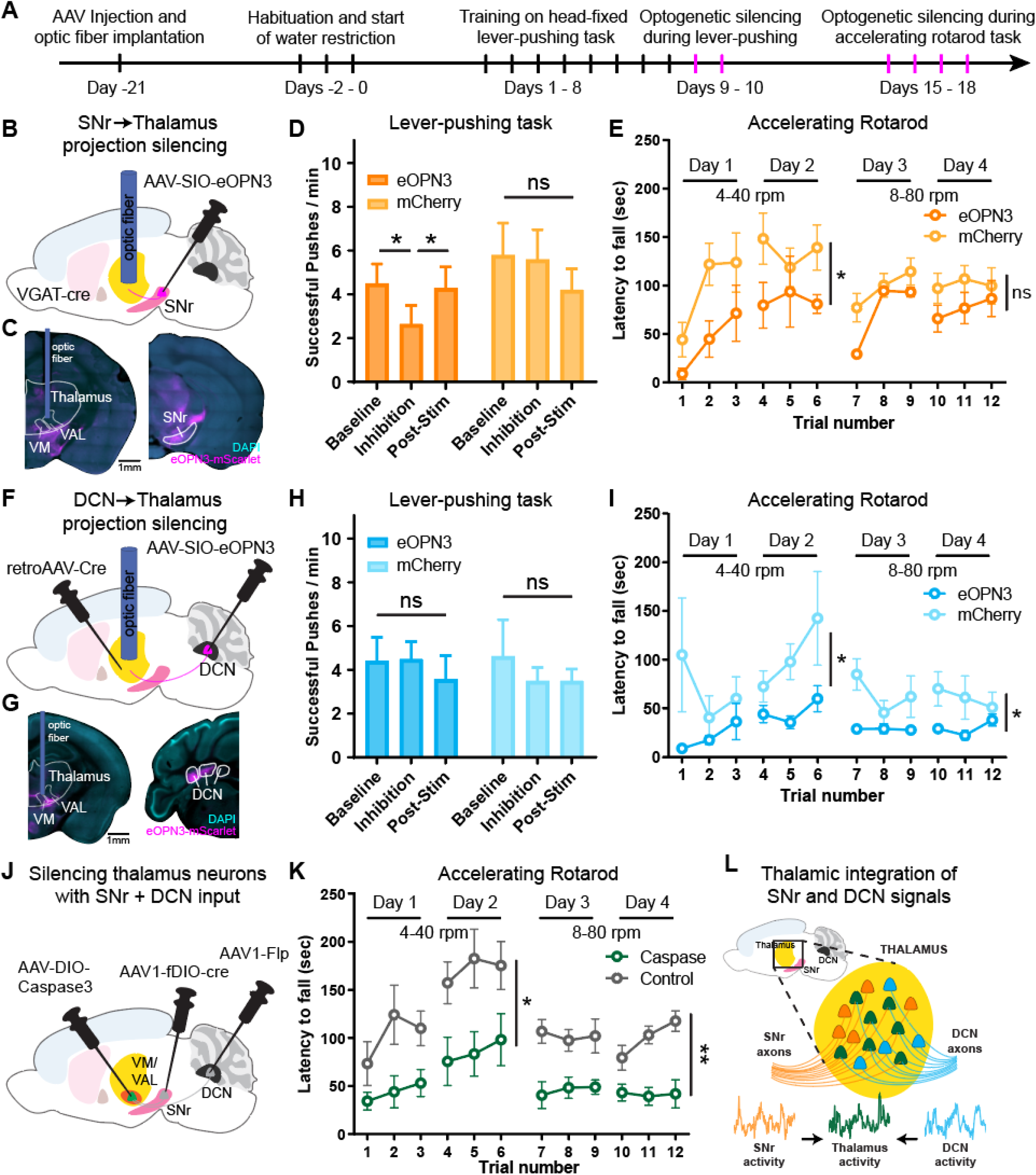
Silencing SNr and DCN inputs to thalamus and dual-input thalamus neurons impairs motor behaviors. (A) Timeline of surgeries and behavior experiments for optogenetic silencing of SNr and DCN projections to the thalamus. Optogenetic silencing sessions are labeled in magenta. (B) Schematic diagram of virus injections and optic fiber placement for optogenetic inhibition of SNr projections to thalamus. (C) Example coronal sections showing AAV-eOPN3 (magenta) expression and optic fiber placement. Scale bar, 1 mm. (D) Lever-push performance of mice expressing eOPN3 in SNr (dark orange) and control mice expressing mCherry (light orange). Number of successful lever pushes shown before (baseline), during (inhibition), and after (post-stim) optogenetic silencing of SNr to thalamus projections. Silencing was performed in mice well-trained on the head-fixed lever-pushing task. Bars denote mean. Error bars, SEM. eOPN3: n=7 mice; mCherry: n=5 mice. Two-way ANOVA with Holm-Sidak multiple comparisons test. (E) Performance on an accelerating rotarod task of mice expressing eOPN3 in SNr (dark orange) and control mice expressing mCherry (light orange). SNr to thalamus projections were unilaterally silenced during each training session. Circles denote mean. Error bars, SEM. eOPN3: n=4 mice; mCherry: n=7 mice. Two-way ANOVA, separately for days 1-2 and days 3-4. (F) Schematic diagram of virus injections and optic fiber placement for optogenetic inhibition of DCN projections to thalamus. (G) Example coronal sections showing AAV-eOPN3 (magenta) expression and optic fiber placement. Scale bar, 1 mm (H) Lever-push performance of mice expressing eOPN3 in DCN (dark blue) and control mice expressing mCherry (light blue). Number of successful lever pushes shown before (baseline), during (inhibition), and after (post-stim) optogenetic silencing of DCN to thalamus projections. Silencing was performed in mice well-trained on the head-fixed lever-pushing task. Bars denote mean. Error bars, SEM. eOPN3: n=8 mice; mCherry: n=7 mice. Two-way ANOVA with Holm-Sidak multiple comparisons test. (I) Performance on an accelerating rotarod task of mice expressing eOPN3 in DCN (dark blue) and control mice expressing mCherry (light blue). DCN to thalamus projections were bilaterally silenced during each training session. Circles denote mean. Error bars, SEM. eOPN3: n=10 mice; mCherry: n=8 mice. Two-way ANOVA, separately for days 1-2 and days 3-4. (J) Schematic diagram of virus injections for ablating thalamic neurons receiving SNr and DCN inputs using Caspase3. (K) Performance on an accelerating rotarod task of mice with bilaterally ablated thalamic neurons receiving SNr and DCN inputs (green) and control mice injected with Caspase3 but no Flp and Cre (gray). Circles denote mean. Error bars, SEM. Caspase: n=6 mice; Control: n=7 mice. Two-way ANOVA, separately for days 1-2 and days 3-4. (L) Proposed model with a fraction of neurons in thalamus (mainly VM and VAL) receiving inputs from SNr and DCN. Neuronal activity from SNr and DCN is integrated in these thalamic neurons and their activity is critical for proper motor learning and execution. ns: not significant, ^∗^p<0.05, ^∗∗^p<0.01

We next assessed locomotion behavior by training these mice on an accelerating rotarod task and inhibiting SNr or DCN projections over the course of learning (**Fig. S4G**). These projections were specifically inhibited during each training session when the mouse was on the rotarod. While control mice learned to stay on the rotarod for longer before falling across two training days at lower rotation speeds and two training days at higher speeds, mice with inhibited SNr inputs had shorter latencies to fall, particularly at slower rotation speeds (**Fig. 4E**). Inhibiting DCN inputs impaired behavioral performance on the rotarod task at both slow and fast speeds (**Fig. 4I**). Interestingly, this deficit was specific to bilateral inhibition of DCN projections to the thalamus, whereas unilateral inhibition did not have any effect (**Fig. S4H**).

Finally, to directly test the causal role of thalamic neurons receiving converging SNr and DCN inputs in motor behavior, we ablated these cells by expressing Caspase3 using the transsynaptic AAV1 approach with Cre and Flp (**Fig. 4J**). In the accelerating rotarod task, we found that while control mice injected with Caspase3 alone learned to stay on the rod, mice with bilateral thalamic neuron ablation were significantly impaired (**Fig. 4K**). Together, these data suggest that SNr and DCN projections to the thalamus and thalamic neurons receiving converging inputs are critical for learning and executing motor tasks.

## Discussion

In this study, we identified a population of neurons in the motor thalamus that receive direct converging synaptic input from the BG and CB (**Fig. 1**). Our electrophysiological and viral tracing data demonstrate that BG and CB projections in the thalamus do not remain separate, instead converge onto individual thalamic neurons, particularly concentrated at the VM and VAL border (**Fig. 2**). This anatomical convergence places these neurons to functionally integrate BG and CB motor signals. Using multi-site fiber photometry, we found that the activity of neurons with SNr and DCN inputs in the thalamus can be predicted from SNr and DCN activity. Importantly, with motor learning, movement-aligned activity within these two inputs and the dual-input cells adapts distinctly in the thalamus (**Fig. 3**), suggesting that the thalamus is not just passively relaying BG and CB activity to the cortex but can act as an active integrator and modulator of these input signals. Indeed, silencing SNr or DCN projections in the thalamus impairs motor behaviors in accordance with their a task-specific activity, and ablating dual-input thalamic neurons severely impairs motor learning (**Fig. 4**). Together, our data reveal a novel role for the thalamus in anatomically and functionally integrating BG and CB motor signals (**Fig. 4L**).

By mapping the location of neurons with BG and CB input to published brain atlases, we provide a valuable resource for the anatomical convergence of these two key motor circuits. We found that their highest density is in the VM and VAL, with a particular enrichment at the border region between these two nuclei of the motor thalamus. We also identified a significant number of neurons with BG and CB inputs in intralaminar regions of the thalamus. Together, these are thalamic regions that have been previously reported to receive axonal projections from BG, CB, or both, consistent with the idea that thalamic integration of BG and CB signals might occur at border regions between the distinct BG and CB recipient areas of the thalamus^22^. An interesting open question is whether these neurons have unique projection patterns or molecular profiles, as reported for neurons with either BG or CB input^7,55^.

Our experiments recording neuronal population activity showed that different motor tasks can differentially engage BG or CB circuits, wherein a lever-pushing task primarily increased SNr activity and treadmill running increased both SNr and DCN activity. Notably, despite SNr projections being inhibitory, their downstream target neurons in the thalamus are activated during both motor tasks. Although SNr activity is generally considered movement suppressive, increased activity of SNr neurons during motor behavior has been previously shown^45,56,57^. The counter-intuitive excitation of SNr target neurons in the thalamus might arise through an entrainment mechanism, such as seen within the CB where synchronous activity of inhibitory Purkinje cells leads to increased downstream activity of DCN neurons^58^. Alternatively, thalamic neurons could also be driven by rebound low-threshold firing bursts following transient pauses in SNr activity^59–61^. Further studies using electrophysiological recordings or high-speed voltage imaging would help distinguish these mechanisms and reveal how these properties support BG and CB signal integration in the thalamus.

Although both SNr and DCN activity increased on average during movement, each of these regions comprise diverse neuronal subpopulations with distinct activity and projection profiles^4,38,45,62–64^. It is thus likely that only specific subpopulations of SNr and DCN neurons converge in the thalamus and contribute to the functional integration we observed during motor learning. Future studies recording activity with cellular resolution in these brain regions will be necessary to further delineate this circuit convergence.

Our finding of functional integration of BG and CB circuits in the thalamus may also have implications for understanding the etiology of movement disorders involving deficits in either the BG or CB systems, such as Parkinson’s disease and ataxia, respectively. Understanding how these two motor systems interact can reveal how one system could potentially compensate for deficits caused by disease in the other.

## Acknowledgments

We thank Dr. W. Xin and members of the Ding lab for insightful discussions and thoughtful comments. This research was funded by NIH/NINDS K99NS130078 (R.H.R.), R01NS081071 (T.M.), RF1NS133599 (T.M.), R01NS091144 (J.B.D.), a Parkinson’s Foundation Postdoctoral Fellowship (R.H.R.), Aligning Science Across Parkinson’s (ASAP-020551) through the Michael J. Fox Foundation for Parkinson’s Research (MJFF) (J.B.D.), a Catalyst grant from the Phil & Penny Knight Initiative for Brain Resilience at the Wu Tsai Neurosciences Institute, Stanford University, and GG gift fund (J.B.D.). For the purpose of open access, the author has applied a CC BY public copyright license to all author accepted manuscripts arising from this submission.

## Author contributions

R.H.R. and J.B.D. conceived the project and designed experiments. R.H.R. performed viral tracing, behavioral, fiber photometry, and silencing experiments. M.A.M. and T.M. performed the registration to published atlases. C.J.H. and R.H.R designed and analyzed behavior and fiber photometry experiments. F.-J.H. performed and R.H.R analyzed slice electrophysiology experiments. Y.S. and C.M. performed cell quantification analysis. R.H.R. and J.B.D. wrote the manuscript with input from all authors.

## Inclusion and diversity

We worked to ensure sex balance in the selection of non-human subjects. One or more of the authors of this paper self-identifies as a member of the LGBTQ+ community. While citing references scientifically relevant for this work, we also actively worked to promote gender balance in our reference list.

## Methods

### Key resources table

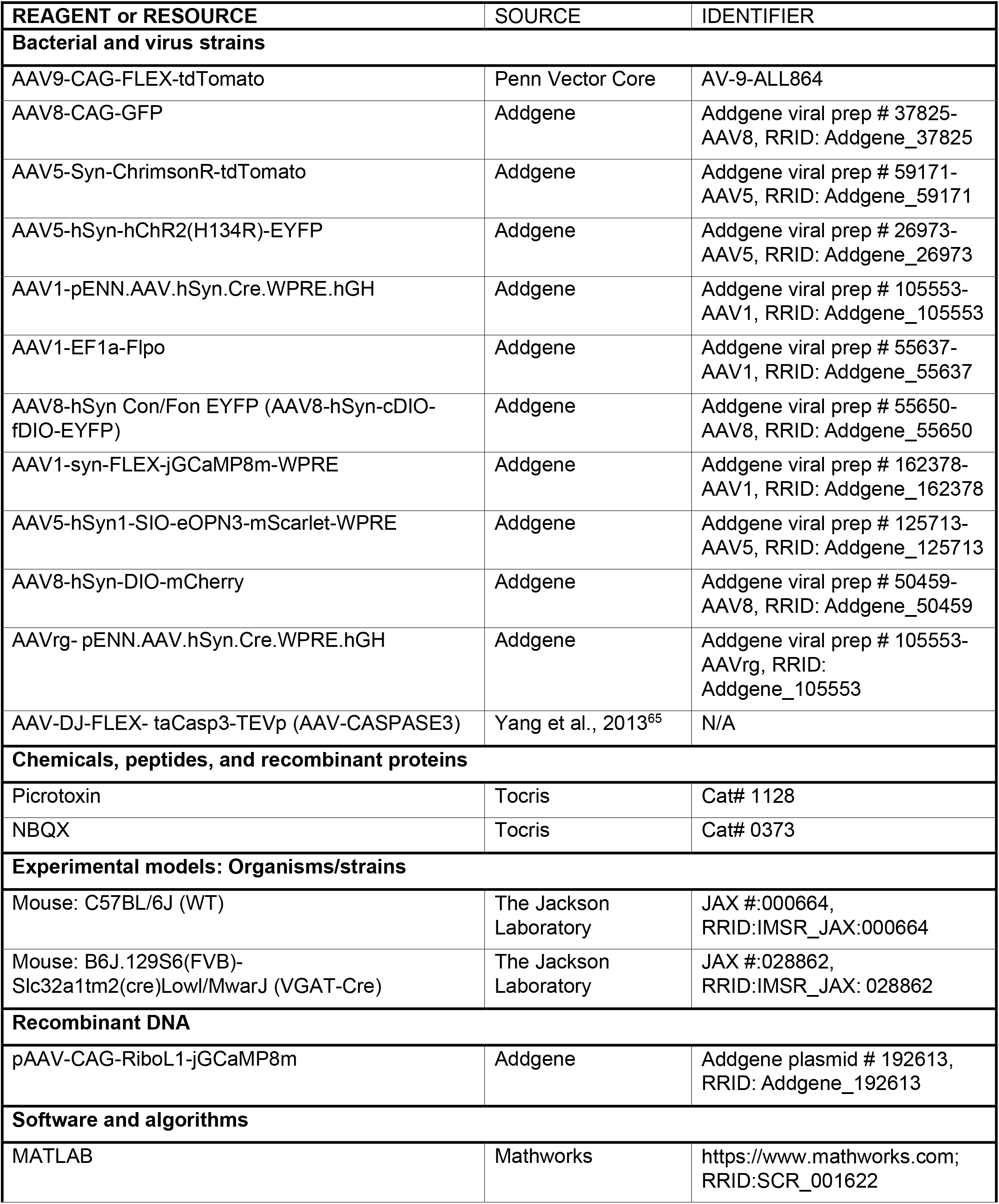
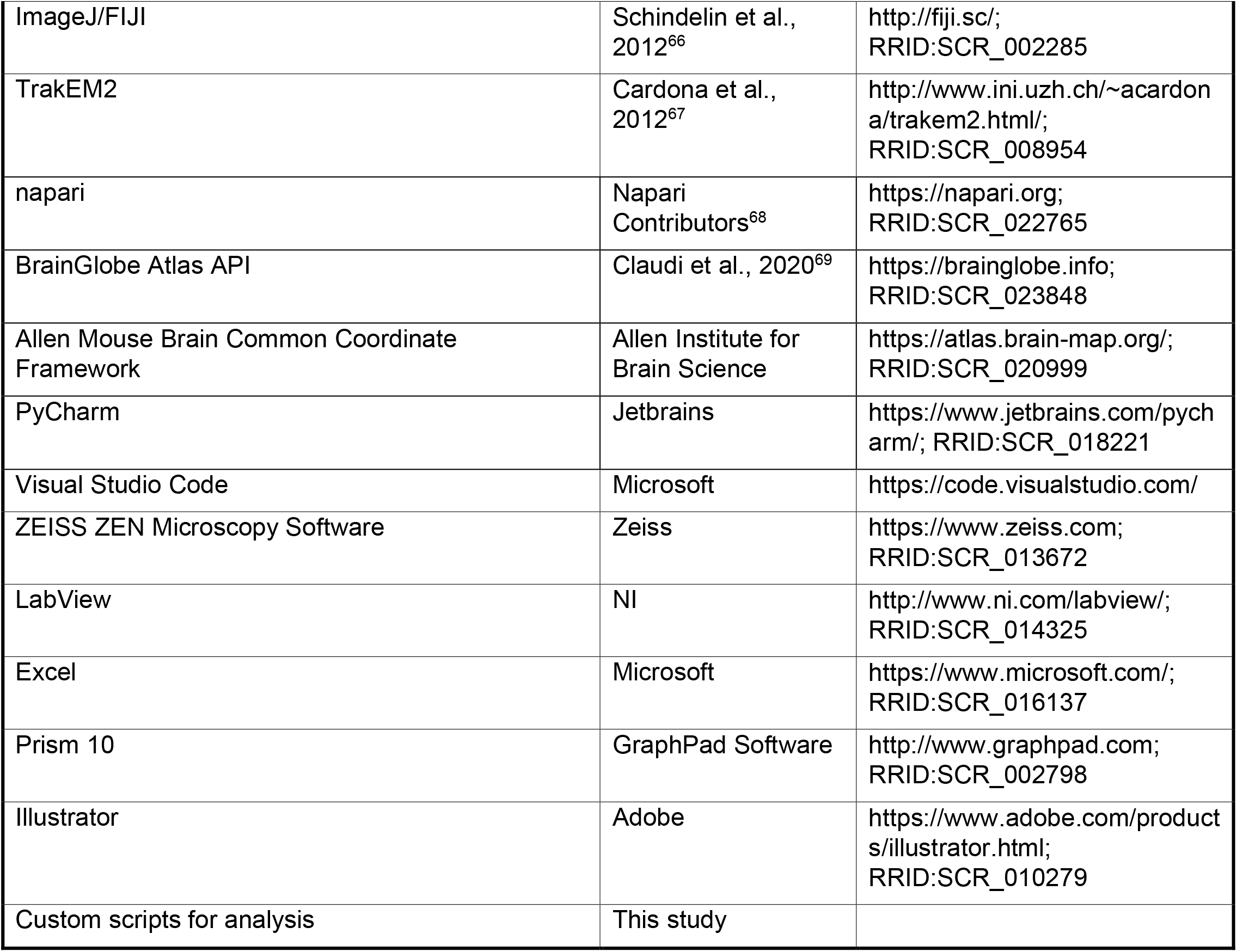

## Resource availability

### Lead contact

Further information and requests for resources and reagents should be directed to and will be fulfilled by the Lead Contact: Jun B. Ding.

### Materials availability

This study did not generate new unique reagents.

### Data and code availability

All new data and code generated in this study are available in the manuscript or will be available on Zenodo by publication. All protocols and key lab materials used and generated in this study are listed and will be available by publication. Any additional information required to reanalyze the data reported in this paper is available from the lead contact upon request.

## Experimental model and subject details

### Animals

All experiments were performed in accordance with protocols approved by the Stanford University Animal Care and Use Committee in keeping with the National Institutes of Health’s Guide for the Care and Use of Laboratory Animals. All mice were kept on a 12 h:12 h light/dark cycle. For behavior and in vivo fiber photometry experiments mice were kept in a reversed light cycle room allowing experiments to be performed during the animals’ wake phase. Mice were habituated to the reversed light cycle for at least two weeks before experiments. All behavior experiments were carried out at consistent times of day to avoid circadian and sleep-related effects. Both male and female mice were used for all experiments. All experiments were performed with adult mice at 2-4 months of age. Mouse lines used in this study include VGAT-Cre (JAX 028862) and Wildtype (WT) C57BL/6J (JAX 000664) mice, obtained from The Jackson Laboratory. Whenever possible, mice were group-housed over the course of experiments.

## Method details

### Stereotaxic viral injection

Stereotaxic injections were performed in 2-4 months old mice under isoflurane anesthesia. For SNr injections, 300-350 nl (60 nl for axonal projection tracing) of virus was slowly injected into the left hemisphere at anterior-posterior (AP) 2.95 mm, medial-lateral (ML) 1.5 mm, and dorsal-ventral (DV) 4.5 mm from bregma. For DCN injections, two injections of 250 nl each were made in the right hemisphere at AP 2.0 mm, ML 2.0 mm and 1.5 mm, and DV 2.25 mm from lambda. For thalamus injections, either three injections of 300 nl each were made in the left hemisphere at AP 1.3 mm, ML 0.8 mm, and DV 4.0 mm, AP 1.3 mm, ML 1.2 mm, and DV 4.0 mm, and AP 1.7 mm, ML 1.0 mm, and DV 3.9 mm from bregma for reporter virus to spread across the entire ventral thalamus, or a single injection of 350-400 nl was made at AP 1.4 mm, ML 1.0 mm, and DV 4.0-4.3 mm from bregma. For experiments using bilateral labeling, same injection volumes and coordinates were used for the contralateral hemisphere. AAV8-hSyn-cDIO-fDIO-EYFP and AAV9-CAG-FLEX-tdTomato virus was used at a titer of ∼1×10^12^ vg/mL, all other viruses were used at a titer ≥ 1×10^13^ vg/mL. For all experiments involving both SNr and DCN injections, such as Cre and Flp expression or ChR2 and ChrimsonR expression, the virus combinations were switched in half of the mice per experimental group, such that, for example, the same number of mice were injected with AAV1-Cre in SNr and AAV1-Flp in DCN as were injected with AAV1-Flp in SNr and AAV1-Cre in DCN. After injection, the scalp was sutured, and the mice were returned to their housing cages. Animals were allowed to recover from surgery for 2-3 weeks before behavior experiments.

### Brain slice histology

Mice were anesthetized with isoflurane and transcardially perfused with phosphate-buffered saline (PBS) and 4% paraformaldehyde (PFA). The brain was dissected out and post-fixed in 4% PFA/PBS for an additional 2 h at room temperature. The brain was transferred to a 30% sucrose/PBS solution for at least 48 h and frozen and sectioned coronally using a sliding microtome (Leica). For mapping cells to published atlases, 40 μm thick slices of the whole brain were cut and mounted in DAPI containing mounting media (VectaShield H-1500, Vector Laboratories). Multi-channel tiled images were obtained using a widefield microscope (Axio Imager, Zeiss) using a 5x objective and a pixel size of 0.908 μm/px or a 10x objective and a pixel size of 0.454 μm/px. Multi-channel images were further processed by linear unmixing. For all other images, 100 μm thick slices of the whole brain were cut and mounted in DAPI containing mounting media (VectaShield H-1500, Vector Laboratories). Multi-channel tiled images were obtained using a laser scanning confocal microscope (LSM900, Zeiss). Representative images shown in figures were median filtered and contrast enhanced.

### Whole-cell slice electrophysiology

Mice (both male and female) were anesthetized and decapitated according to an animal protocol approved by the Stanford University Animal Care and Use Committee. The extracted brains were submerged in ice-cold artificial cerebrospinal fluid (ACSF) containing (in mM) 125 NaCl, 2.5 KCl, 1.25 NaH_2_PO_4_, 25 NaHCO_3_, 15 glucose, 2 CaCl_2_ and 1 MgCl_2_, with continuous bubbling with 95% O_2_ and 5% CO_2_ (300-305 mOsm, pH 7.4). Coronal slices (300 μm thickness) containing thalamus were prepared with a vibratome (Leica). Slices were then incubated for 30 min in 34°C ACSF for recovery and then held at room temperature. For patch-clamp recordings, slices were transferred to a recording chamber, held in place with an anchor (Warner Instrument), and mounted onto a microscope (BX51, Olympus). The slices were continuously perfused with ACSF at a rate of 2-3 ml/min at 30°C during recording. Patch-clamp recordings were made through a Multiclamp 700B amplifier (Molecular Devices) and monitored with custom-made MATLAB (Mathworks) software. The signal was low-pass filtered at 2.2 kHz and digitized at 10 kHz (NI PCIe-6259 card, National Instrument). For whole cell voltage-clamp, glass pipettes (2.5-4.5 MΩ) were filled with internal solution containing (in mM) 115 CsMeSO_3_, 10 TEA, 10 HEPES, 5 QX-314 chloride, 4 Mg-ATP, 0.4 Na_3_-GTP, 10 Na_2_-phosphocreatine, 1 EGTA, 0.1 CaCl_2_ (280-290 mOsm, adjusted to pH 7.3-7.4 with CsOH). For measuring optogenetic-induced EPSCs and IPSCs, the membrane potential was held at -70 mV and +10 mV respectively and single optogenetic stimulation pulses were applied to the whole imaging field (Opto Engine LLC) with 1 ms 1 mW (measured at the front-aperture of the objective) 450 nm pulses for ChR2 activation and 50 ms 10 mW 594 nm pulses for ChrimsonR activation. These wavelengths and laser powers were chosen to reduce cross-activation of ChrimsonR with 450 nm light or ChR2 with 594 nm light. The location of recorded neurons was documented based on low-magnification DIC and fluorescence images taken at the conclusion of each recording. Cells with series resistance >25 MΩ were excluded.

### In vivo fiber photometry

Following stereotaxic virus injection, optic fiber cannulae were implanted in mice under isoflurane anesthesia for multi-site in vivo fiber photometry. Optic fiber cannula with a 200 μm core, 0.39 NA, and 1.25 mm diameter ferrule (RWD Life Science) were implanted at AP 2.95 mm, ML 1.5 mm, and DV 4.2 mm from bregma for SNr, AP 2.0 mm, ML 1.0 mm, and DV 2.0 mm from lambda for DCN, and AP 1.4 mm, ML 1.0 mm, and DV 4.0 mm from bregma for the thalamus. 4.5 mm long optic fiber cannulae were used for SNr and thalamus, 2.5 mm long cannula were used for DCN. The cannulae were fixed to the skull using dental cement (C&B Metabond, Parkell). At the end of surgery, a custom titanium headplate was attached to the skull to fixate the head to the behavior stage during subsequent experiments. Animals were allowed to recover from surgery for 2 weeks before behavior and recording experiments.

For fiber photometry recordings of calcium activity, soma-targeted GCaMP8m (AAV8-CAG-RiboL1-jGCaMP8m, packaged in lab) was mixed 1:1 with either AAV1-Cre or AAV1-Flp and injected in SNr and DCN. AAV1-syn-FLEX-jGCaMP8m-WPRE was expressed in the thalamus. See “Stereotaxic viral injection” for coordinates. We used soma-targeted GCaMP in the SNr and DCN to avoid Ca^2+^ signal from their axonal projections in the thalamus contributing to the thalamus recordings^48^.

Multi-site in vivo fiber photometry recordings were made using a dual color multichannel fiber photometry system (R811, RWD Life Science) and a low-auto fluorescence 1-4 fan out bundled fiber patch cord, with a 200 μm core, 0.39 NA, and a 1.25 mm diameter ferrule. GCaMP8m was stimulated at 470 nm and 410 nm for its isosbestic control and green fluorescence emission was recorded at 30 Hz. In addition to signal from the three brain regions (SNr, DCN, thalamus), an additional background signal on the photometry system camera was recorded. Fiber photometry and behavior recordings were synchronized using a frame trigger output from the photometry system as was as a high-speed IR camera. Only mice performing on the lever-pushing task (average ≥10 lever pushes per sessions) were included for analysis of fiber photometry experiments.

### Head-fixed lever-pushing task

3-4 months-old mice were water restricted and fed with 1 ml of water per day. 2-3 days after the start of water restriction their body weight was reduced to 85-90% of baseline weight and maintained throughout training. Following water restriction mice were habituated to head fixation and received water from a water port for at least three 5–20-minute sessions. Habituated mice were trained for 10 days with one 15-minute training session per day. The training apparatus was a custom-built robotic manipulandum based on the design by Wagner et al., 2020^70^ and set-up so that mice can perform 8 mm linear pushing movements using their right limb. During each training session mice could self-initiate lever pushes. Lever movements that were greater than 1 mm were registered as pushes. Movements that cross the set 5 mm threshold were rewarded with a small drop of water and considered successful.

### Head-fixed locomotion task

Following training on the lever-pushing task, mice were trained on a head-fixed locomotion task. This custom-built training apparatus consisted of a cylindrical rod treadmill with a 16 cm diameter. Mice were trained for 5 days with one 15-minute training session per day. During this time mice were head-fixed on the behavioral apparatus and their position adjusted so that the four limbs were resting on the treadmill. Mice were allowed to run on the treadmill at will. Treadmill movement was measured using a high-speed IR camera, as well as a rotary encoder.

### Accelerating rotarod task

For the accelerating rotarod task, mice were trained for 4 days with three training sessions each. During the first 2 days, the rotarod (AccuRotor EzRod) was set up to accelerate from 4 rpm to 40 rpm linearly over 5 minutes. During the last 2 days, the rotarod was set up to accelerate from 8 rpm to 80 rpm. Each day, mice were placed on the rotarod for three sessions, each 1 h apart. Once mice were placed on the rotarod and were stable on the stationary rod, the motor was turned on. Behavioral performance was quantified by measuring the time it took for the mouse to fall off the rod from the beginning of rotation.

### Optogenetic silencing of projection terminals

Following stereotaxic virus injection, optic fiber cannulae were implanted in mice under isoflurane anesthesia for optogenetic silencing. Optic fiber cannula with a 400 μm core, 0.5 NA, 4.5mm length, and either a 2.5mm diameter ferrule for unilateral silencing or a 1.25 mm diameter ferrule for bilateral silencing (RWD Life Science) were implanted at AP 1.4 mm, ML 1.0 mm, and DV 4.0 mm from bregma targeting the thalamus. The cannulae were fixed to the skull using dental cement (C&B Metabond, Parkell). At the end of surgery, a custom titanium headplate was attached to the skull to fixate the head to the behavior stage during subsequent experiments. Animals were allowed to recover from surgery for 2 weeks before behavior and recording experiments.

For optogenetic silencing of SNr projections to the thalamus, AAV5-hSyn1-SIO-eOPN3-mScarlet-WPRE or AAV8-hSyn-DIO-mCherry (control) was injected in the SNr of VGAT-Cre mice. For optogenetic silencing of DCN projections to the thalamus AAV5-hSyn1-SIO-eOPN3-mScarlet-WPRE or AAV8-hSyn-DIO-mCherry (control) was injected in the DCN, as well as AAVrg-hSyn.Cre.WPRE.hGH in the thalamus. See “Stereotaxic viral injection” for coordinates.

SNr or DCN projections were silenced in mice well-trained (for 8-11 days) on the head-fixed lever-pushing task. During two optogenetic silencing sessions following head-fixed lever-pushing training, mice were first allowed to perform the task during a 2-or 3-minute baseline period without inhibition. eOPN3 or mCherry control expressing SNr or DCN projections were then inhibited for 3 minutes using a 594 nm laser (Opto Engine LLC) with 500 ms pulses at 0.1 Hz and a power of 10 mW from the fiber tip. Lever-push behavior was quantified using the 2 min before optogenetic silencing as baseline, the 2 min window starting 1 min after optogenetic laser onset as inhibition period, and the 2 min window starting 1 min after optogenetic silencing ended as post-stim washout period. Out of the two optogenetic silencing sessions the session with the higher number of lever-pushes during baseline was used for analysis. Only mice with ≥2 lever pushes per minute during the baseline were included.

For silencing SNr and DCN projections during the accelerating rotarod task, eOPN3 or mCherry control was stimulated starting 1 min before mice were placed on the rotarod and throughout each rotarod session. Optogenetic silencing was terminated once mice fell off the rod.

## Quantification and statistical analysis

### Analysis of patch-clamp recordings

The amplitude of optogenetically evoked EPSCs or IPSCs was calculated by subtracting the mean baseline current during the 50 ms before the laser was turned on from the peak current following laser onset. In some cases of mice expressing ChrimsonR in the SNr and ChR2 in the DCN, residual cross-activation of ChrimsonR at 450nm led to an inward current due to the reversed GABA potential at -70mV. By comparing the current amplitude at -70 mV and +10 mV with 450 nm light stimulation, we determined if the -70 mV signal was an actual oEPSC caused by 450 nm stimulation of ChR2 or cross-activation of ChrimsonR. Cells with an inward current ≥ 25 pA when stimulating DCN projections were considered as receiving DCN input, and cells with an outward current ≥ 100pA when stimulating SNr projections were considered as receiving SNr input.

### Quantification and registration of neurons to published atlases

Serial-section widefield coronal images were first level-balanced in FIJI^66^ using Python scripts such that the mean background pixel intensity was equivalent across all sections from each mouse. Only global (section-wide) adjustments were permitted; local manipulations were never made. Adjusted images were downsampled to ∼3.63 μm/px and imported into TrakEM2 software^67^ for serial-section alignment and annotation. Adjacent sections were semi-manually aligned to one another using conspicuous features such as tissue edges and contiguous vasculature.

To programmatically register each reconstructed thalamus to an atlas, we adopted and updated the algorithm from Hunnicutt et al.^7^ (see their Supplementary Table 1) using a combination of fiducial registration points and coronal bounding boxes. Five fiducial points were annotated along the midline: the anterior- and posterior-most edges of the corpus callosum, the posterior edge of the anterior commissure, the anterior emergence of the dentate gyrus, and the position of the subcommissural organ (SCO) just posterior to the end of the habenula. Additionally, two matching pairs of points were annotated along the lateral edges of the thalamus to facilitate corrections for left/right asymmetry in coronal sections: the posterior-most limit of the arch of the stria terminalis and posterior edge of the lateral geniculate nucleus. Next, rectangular coronal bounding boxes were annotated for the left hemisphere on all coronal sections starting with the section just posterior to the anterior commissure and ending with the last section containing the medial geniculate nucleus. The medial and lateral bounds of each box were set to the midline and lateral-most edge of the thalamus, respectively. The dorsal bound of the box was set to the dorsal-most edge of the thalamus and/or midbrain, excepting any further dorsal extension made solely by the habenula (if present) and limited to the midline ventral edge of the sensory-related component of the superior colliculus (e.g., the optic layer; if present). The ventral bound of the box was defined in two parts based on the anterior-most section where the dorsal edge of the optic nerve crosses the ventral edge of the internal capsule. Anterior to this section, the ventral bound is defined by the dorsal edge of the third ventricle. Going posterior from this section, the ventral bound is defined by the bottom edge of the internal capsule and/or cerebral peduncle. Collectively, these bounding boxes defined a 3D registration volume.

Experimental brains were independently registered to both the Allen Mouse Brain Common Coordinate Framework (CCFv3)^29^ and the Franklin-Paxinos-based Unified Anatomical Atlas^30^. Both atlases were accessed at 10 μm resolution using the BrainGlobe Atlas API^69^ within the napari Python environment^68^. For each atlas, coordinates for equivalent fiducial landmarks and coronal bounding boxes as above were programmatically extracted. The atlas volume was then mapped to each experimental brain using a series of affine transformation steps: 1) the best-fit affine solution that maps the first four fiducial landmarks (corpus callosum ×2, anterior commissure, and dentate gyrus) from source to target, only applied to the AP and LM axes; 2) a shear transform along the A-P axis to compensate for left/right asymmetry; 3) a translation to anchor source and target volumes at the SCO fiducial, which becomes the coordinate origin for subsequent steps; 4) a scaling transform on the A-P axis to match 99% of the bounding box volume from its anterior pole to the SCO fiducial; and 5-6) scaling transforms on the LM and DV axes to match the average width and height of individual bounding boxes within 1 mm anterior and 1 mm posterior of the SCO fiducial. Following these steps, a final transformation was applied on a per-slice basis to compensate for any drift in serial section alignment based on the distance between the dorsomedial corner of the bounding box for each slice and the equivalent corner of the bounding box derived from the intersection of the transformed registration volume with the slice plane. By combining these steps, we defined a set of affine transformations that mapped atlas data to each experimental brain section. To register experimental coordinate data to atlas coordinate space, the inverse transformation for each brain slice was used.

eYFP-positive neurons were annotated in widefield images (at 0.454 μm/px) using the Cell Counter FIJI plugin and a graphics tablet (22HD; Wacom). As eYFP-positive neurons exhibited a wide range of intensity levels, the observer was free to dynamically adjust image contrast during the annotation process.

Two additional observers independently annotated a randomized subset of images to verify accuracy and achieved comparable counts and distributions of neurons per slice. To mitigate concerns of off-target labeling, cell counts were subsequently filtered based on relative intensity. Neuron intensity was computed by taking a 25 μm-wide circular ROI at each annotation coordinate and averaging the 20% brightest pixels within this ROI. Neuron intensity values were normalized for each brain, and any neurons with normalized intensity <=5% were excluded from subsequent analyses. Annotated neurons (ranging between 428 and 2516 per mouse) were registered to atlas coordinates in napari using the slice-specific transformations obtained above. Neurons were assigned to atlas brain regions by querying the value of the nearest 10-μm atlas voxel to the registered neuron coordinate. Average neuron distribution with respect to atlas region was computed by counting the total number of neurons assigned to each atlas region divided by the total number of eYFP-positive neurons per mouse. Average 3D neuron density was computed by first calculating the percentage of eYFP-positive neurons per mouse that resided within 100 × 100 × 100 μm bins of registered atlas coordinate space, then averaging these binned values across mice.

### Analysis of Ca^2+^ fiber photometry recordings

Data recorded through fiber photometry was first background-subtracted and filtered using a 4th order Butterworth lowpass filter using a critical frequency of 2 Hz. A least-squares linear fit was then applied to the 410 nm isosbestic signal to align it to the 470 nm signal. This fitted 410 nm signal was then subtracted from the 470 nm signal and the result divided by the fitted 410 nm signal, for normalization and controlling for bleaching and movement artifacts.

For linear regression, random 135-second intervals across all recording sessions were chosen for each mouse and a multiple linear regression using A × Ca^2+^_SNr_ + B × Ca^2+^_DCN_ + C = Ca^2+^_Thalamus_ was performed between the recorded Ca^2+^ signal from SNr and DCN and the recorded Ca^2+^ signal from thalamus. For each random interval a predicted thalamus signal was calculated using the coefficients from the linear regression. To test the goodness of fit of this prediction an R^2^ value was calculated for each interval. As a control, the same regression and prediction was performed using data in which the SNr and DCN signals for each interval were shifted in time by a random amount between 0 and ±67.5 sec, each, relative to the thalamus signal. Again, a regression fit (R^2^) was calculated for these predictions for each shuffled interval. All recording intervals were then averaged per mouse.

For aligning Ca^2+^ signals to movement onset, the movement onset for each trial during the lever-pushing and locomotion tasks was calculated based on the forward movement velocity. Recordings were trial segmented for 10 seconds around movement onset and signal pre-processing steps (filtering and normalizing) applied to these trial windows. For each trial a z-score was calculated using the mean and standard deviation of a 700 ms pre-movement baseline period. This baseline period was set immediately before movement onset for the lever-pushing task and shifted by 500 ms prior to movement onset for the locomotion task. This was done to account for changes in Ca^2+^ signals prior to our defined forward movement onset. All trials from early learning sessions (days 1-3 for lever pushing and days 1-2 for locomotion) as well as all trials from late learning sessions (days 8-10 for lever pushing and days 4-5 for locomotion) were then averaged per mouse.

### Statistics

Data analysis was performed in Excel (Microsoft), Matlab (MathWorks), Python, and ImageJ. Statistical analysis was performed in Prism 10 (GraphPad Software). For ANOVA tests we report the main group effect or interaction effect, as stated in figure legends. If significant we also report post-hoc multiple comparison tests as stated in figure legends. All data are presented as mean ± SEM (standard error of the mean) in figures and statistical tests as well as sample sizes noted in figure legends. Statistical thresholds used: ∗ p < 0.05, ∗∗ p < 0.01, ∗∗∗ p < 0.001, ns: non-significant.

## References

1. Klaus, A., Alves da Silva, J. & Costa, R. M. What, If, and When to Move: Basal Ganglia Circuits and Self-Paced Action Initiation. Annu. Rev. Neurosci. 42, annurev-neuro-072116-031033 (2019).

2. Graybiel, A. M., Aosaki, T., Flaherty, A. W. & Kimura, M. The Basal Ganglia and Adaptive Motor Control. Science (80-.). 265, 1826–1831 (1994).

3. Manto, M. et al. Consensus Paper: Roles of the Cerebellum in Motor Control—The Diversity of Ideas on Cerebellar Involvement in Movement. The Cerebellum 11, 457–487 (2012).

4. Carey, M. R. The cerebellum. Curr. Biol. 34, R7–R11 (2024).

5. De Zeeuw, C. I. & Ten Brinke, M. M. Motor Learning and the Cerebellum. Cold Spring Harb. Perspect. Biol. 7, a021683 (2015).

6. Graybiel, A. M. The basal ganglia: learning new tricks and loving it. Curr. Opin. Neurobiol. 15, 638–644 (2005).

7. Hunnicutt, B. J. et al. A comprehensive thalamocortical projection map at the mesoscopic level. Nat. Neurosci. 17, 1276–1285 (2014).

8. Hoover, J. E. & Strick, P. L. The Organization of Cerebellar and Basal Ganglia Outputs to Primary Motor Cortex as Revealed by Retrograde Transneuronal Transport of Herpes Simplex Virus Type 1. J. Neurosci. 19, 1446–1463 (1999).

9. Doya, K. Complementary roles of basal ganglia and cerebellum in learning and motor control. Curr. Opin. Neurobiol. 10, 732–739 (2000).

10. Percheron, G., François, C., Talbi, B., Yelnik, J. & Fénelon, G. The primate motor thalamus. Brain Res. Rev. 22, 93–181 (1996).

11. Haber, S. N. & Calzavara, R. The cortico-basal ganglia integrative network: The role of the thalamus. Brain Res. Bull. 78, 69–74 (2009).

12. McFarland, N. R. & Haber, S. N. Thalamic Relay Nuclei of the Basal Ganglia Form Both Reciprocal and Nonreciprocal Cortical Connections, Linking Multiple Frontal Cortical Areas. J. Neurosci. 22, 8117–8132 (2002).

13. Nambu, A., Yoshida, S. & Jinnai, K. Projection on the motor cortex of thalamic neurons with pallidal input in the monkey. Exp. Brain Res. 71, 658–662 (1988).

14. Herkenham, M. The afferent and efferent connections of the ventromedial thalamic nucleus in the rat. J. Comp. Neurol. 183, 487–517 (1979).

15. Alexander, G. E. & Crutcher, M. D. Functional architecture of basal ganglia circuits: neural substrates of parallel processing. Trends Neurosci. 13, 266–271 (1990).

16. Nakamura, K. C., Sharott, A. & Magill, P. J. Temporal Coupling with Cortex Distinguishes Spontaneous Neuronal Activities in Identified Basal Ganglia-Recipient and Cerebellar-Recipient Zones of the Motor Thalamus. Cereb. Cortex 24, 81–97 (2014).

17. Kuramoto, E. et al. Complementary distribution of glutamatergic cerebellar and GABAergic basal ganglia afferents to the rat motor thalamic nuclei. Eur. J. Neurosci. 33, 95–109 (2011).

18. Bosch-Bouju, C., Hyland, B. I. & Parr-Brownlie, L. C. Motor thalamus integration of cortical, cerebellar and basal ganglia information: implications for normal and parkinsonian conditions. Front. Comput. Neurosci. 7, 163 (2013).

19. Middleton, F. A. & Strick, P. L. Basal ganglia and cerebellar loops: motor and cognitive circuits. Brain Res. Rev. 31, 236–250 (2000).

20. Hoshi, E., Tremblay, L., Féger, J., Carras, P. L. & Strick, P. L. The cerebellum communicates with the basal ganglia. Nat. Neurosci. 8, 1491–1493 (2005).

21. Bostan, A. C., Dum, R. P. & Strick, P. L. The basal ganglia communicate with the cerebellum. Proc. Natl. Acad. Sci. U. S. A. 107, 8452–6 (2010).

22. Hintzen, A., Pelzer, E. A. & Tittgemeyer, M. Thalamic interactions of cerebellum and basal ganglia. Brain Struct. Funct. 223, 569–587 (2018).

23. Koster, K. P. & Sherman, S. M. Convergence of inputs from the basal ganglia with layer 5 of motor cortex and cerebellum in mouse motor thalamus. Elife 13, (2024).

24. Buee, J., Deniau, J. M. & Chevalier, G. Nigral modulation of cerebello-thalamo-cortical transmission in the ventral medial thalamic nucleus. Exp. Brain Res. 65, 241–244 (1986).

25. Sakai, S. T., Stepniewska, I., Qi, H. X., Kaas, J. H. & Sakai, S. T. Pallidal and Cerebellar Afferents to Pre-Supplementary Motor Area Thalamocortical Neurons in the Owl Monkey: A Multiple Labeling Study. J. Comp. Neurol. 417, 164–180 (2000).

26. Deniau, J. M., Kita, H. & Kitai, S. T. Patterns of termination of cerebellar and basal ganglia efferents in the rat thalamus. Strictly segregated and partly overlapping projections. Neurosci. Lett. 144, 202–206 (1992).

27. Sakai, S. T. & Patton, K. Distribution of cerebellothalamic and nigrothalamic projections in the dog: A double anterograde tracing study. J. Comp. Neurol. 330, 183–194 (1993).

28. Anderson, M. E. & DeVito, J. L. An analysis of potentially converging inputs to the rostral ventral thalamic nuclei of the cat. Exp. Brain Res. 68, (1987).

29. Wang, Q. et al. The Allen Mouse Brain Common Coordinate Framework: A 3D Reference Atlas. Cell 181, 936-953.e20 (2020).

30. Chon, U., Vanselow, D. J., Cheng, K. C. & Kim, Y. Enhanced and unified anatomical labeling for a common mouse brain atlas. Nat. Commun. 10, 5067 (2019).

31. Franklin, K. & Paxinos, G. The Mouse Brain in Stereotaxic Coordinates, Compact -3rd Edition. Academic Press (2008).

32. Heck, D. H., Fox, M. B., Correia Chapman, B., McAfee, S. S. & Liu, Y. Cerebellar control of thalamocortical circuits for cognitive function: A review of pathways and a proposed mechanism. Frontiers in Systems Neuroscience vol. 17 (2023).

33. Gao, Z. et al. A cortico-cerebellar loop for motor planning. Nature 563, (2018).

34. Judd, E. N., Lewis, S. M. & Person, A. L. Diverse inhibitory projections from the cerebellar interposed nucleus. Elife 10, (2021).

35. Kebschull, J. M. et al. Cerebellar nuclei evolved by repeatedly duplicating a conserved cell-type set. Science (80-.). 370, (2020).

36. Gornati, S. V. et al. Differentiating Cerebellar Impact on Thalamic Nuclei. Cell Rep. 23, 2690–2704 (2018).

37. Strick, P. L., Dum, R. P. & Fiez, J. A. Cerebellum and nonmotor function. Annual Review of Neuroscience vol. 32 (2009).

38. McElvain, L. E. et al. Specific populations of basal ganglia output neurons target distinct brain stem areas while collateralizing throughout the diencephalon. Neuron 109, (2021).

39. Zingg, B. et al. AAV-Mediated Anterograde Transsynaptic Tagging: Mapping Corticocollicular Input-Defined Neural Pathways for Defense Behaviors. Neuron 93, 33–47 (2017).

40. Fenno, L. E. et al. Targeting cells with single vectors using multiple-feature Boolean logic. Nat. Methods 11, 763–772 (2014).

41. Jeljeli, M., Strazielle, C., Caston, J. & Lalonde, R. Effects of ventrolateral-ventromedial thalamic lesions on motor coordination and spatial orientation in rats. Neurosci. Res. 47, 309–316 (2003).

42. Canavan, A. G. M., Nixon, P. D. & Passingham, R. E. Motor learning in monkeys (Macaca fascicularis) with lesions in motor thalamus. Exp. Brain Res. 77, (1989).

43. Dacre, J. et al. A cerebellar-thalamocortical pathway drives behavioral context-dependent movement initiation. Neuron 109, 2326-2338.e8 (2021).

44. Takahashi, N. et al. Thalamic input to motor cortex facilitates goal-directed action initiation. Curr. Biol. 31, 4148-4155.e4 (2021).

45. Jin, X., Tecuapetla, F. & Costa, R. M. Basal ganglia subcircuits distinctively encode the parsing and concatenation of action sequences. Nat. Neurosci. 17, 423–430 (2014).

46. Roth, R. H. & Ding, J. B. Cortico-basal ganglia plasticity in motor learning. Neuron (2024) doi:10.1016/j.neuron.2024.06.014.

47. Sibener, L. J. et al. Dissociable roles of thalamic nuclei in the refinement of reaches to spatial targets. bioRxiv Prepr. Serv. Biol. (2023) doi:10.1101/2023.09.20.558560.

48. Legaria, A. A. et al. Fiber photometry in striatum reflects primarily nonsomatic changes in calcium. Nat. Neurosci. 25, 1124–1128 (2022).

49. Sarnaik, R. & Raman, I. M. Control of voluntary and optogenetically perturbed locomotion by spike rate and timing of neurons of the mouse cerebellar nuclei. Elife 7, (2018).

50. Sheng, M.-J., Lu, D.Shen, Z.-M. & Poo, M.-M. Emergence of stable striatal D1R and D2R neuronal ensembles with distinct firing sequence during motor learning. Proc. Natl. Acad. Sci. U. S. A. 116, 11038–11047 (2019).

51. Becker, M. I. & Person, A. L. Cerebellar Control of Reach Kinematics for Endpoint Precision. Neuron 103, 335-348.e5 (2019).

52. Wagner, M. J., Kim, T. H., Savall, J., Schnitzer, M. J. & Luo, L. Cerebellar granule cells encode the expectation of reward. Nature 544, 96–100 (2017).

53. Ma, L. et al. Locomotion activates PKA through dopamine and adenosine in striatal neurons. Nature 611, (2022).

54. Mahn, M. et al. Efficient optogenetic silencing of neurotransmitter release with a mosquito rhodopsin. Neuron 109, 1621-1635.e8 (2021).

55. Phillips, J. W. et al. A repeated molecular architecture across thalamic pathways. Nat. Neurosci. 22, 1925–1935 (2019).

56. Gulley, J. M., Kuwajima, M., Mayhill, E. & Rebec, G. V. Behavior-related changes in the activity of substantia nigra pars reticulata neurons in freely moving rats. Brain Res. 845, 68–76 (1999).

57. Liu, D. et al. A common hub for sleep and motor control in the substantia nigra. Science (80-.). 367, 440–445 (2020).

58. Person, A. L. & Raman, I. M. Purkinje neuron synchrony elicits time-locked spiking in the cerebellar nuclei. Nature 481, 502–505 (2012).

59. Llinás, R. & Jahnsen, H. Electrophysiology of mammalian thalamic neurones in vitro. Nature 297, 406–8 (1982).

60. Jahnsen, H. & Llinás, R. Electrophysiological properties of guinea-pig thalamic neurones: an in vitro study. J. Physiol. 349, 205–226 (1984).

61. Kim, J. et al. Inhibitory Basal Ganglia Inputs Induce Excitatory Motor Signals in the Thalamus. Neuron 95, 1181-1196.e8 (2017).

62. Lee, J., Wang, W. & Sabatini, B. L. Anatomically segregated basal ganglia pathways allow parallel behavioral modulation. Nat. Neurosci. 23, 1388–1398 (2020).

63. Foster, N. N. et al. The mouse cortico–basal ganglia–thalamic network. 598, 188–194 (2021).

64. Zhu, J., Hasanbegoviż, H., Liu, L. D., Gao, Z. & Li, N. Activity map of a cortico-cerebellar loop underlying motor planning. Nat. Neurosci. 26, 1916–1928 (2023).

65. Yang, C. F. et al. Sexually Dimorphic Neurons in the Ventromedial Hypothalamus Govern Mating in Both Sexes and Aggression in Males. Cell 153, 896–909 (2013).

66. Schindelin, J. et al. Fiji: an open-source platform for biological-image analysis. Nat. Methods 9, 676–682 (2012).

67. Cardona, A. et al. TrakEM2 Software for Neural Circuit Reconstruction. PLoS One 7, e38011 (2012).

68. NapariContributors. napari: a multi-dimensional image viewer for python. (2019) doi:10.5281/zenodo.3555620.

69. Claudi, F. et al. BrainGlobe Atlas API: a common interface for neuroanatomical atlases. J. Open Source Softw. 5, 2668 (2020).

70. Wagner, M. J., Savall, J., Kim, T. H., Schnitzer, M. J. & Luo, L. Skilled reaching tasks for head-fixed mice using a robotic manipulandum. Nat. Protoc. (2020) doi:10.1038/s41596-019-0286-8.

