## Supplemental Figures for "Thalamic integration of basal ganglia and cerebellar circuits during motor learning"

**Roth et al., bioRxiv (2024)**

**Supplemental Figures**

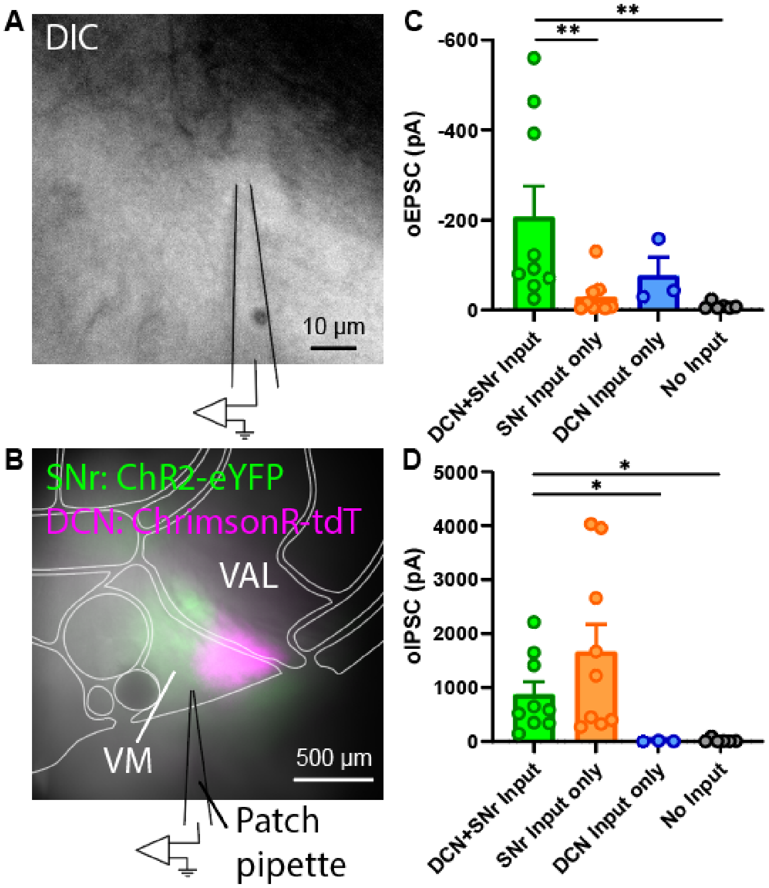


**Roth et al. Figure S1**

**Figure S1 (related to Figure 1): Pathway specific optogenetic stimulation and whole-cell patch clamp recordings in thalamus.**

(A) Example DIC image of neuron recorded in thalamus. Scale bar, 10 μm.
(B) Example DIC and fluorescence image showing location of recorded neuron in VM thalamus. Axonal projections from SNr express ChR2-eYFP (green) and projections from DCN express ChrimsonR-tdTomato (red). Scale bar, 500 μm.
(C) oEPSC amplitudes in response to optogenetic stimulation of DCN projections. Bars denote mean and circles individual cells. Error bars, SEM. DCN+SNr inputs: -206.6±68.4 pA, n=9 cells; SNr input only: -28.6±13.7 pA, n=9 cells; DCN input only: -76.9±40.9 pA, n=3 cells; No input: -8.9±3.1 pA, n=6 cells. Kruskal-Wallis test with Dunn's multiple comparisons test.
(D) oIPSC amplitudes in response to optogenetic stimulation of SNr projections. Data in C and D are from a mix of mice expressing ChrimsonR in SNr and ChR2 in DCN or vice versa. Bars denote mean and circles individual cells. Error bars, SEM. DCN+SNr inputs: 874.9±236.5 pA, n=9 cells; SNr input only: 1666.6±511.5 pA, n=9 cells; DCN input only: 9.2±5.7 pA, n=3 cells; No input: 21.3±13.7 pA, n=6 cells. Kruskal-Wallis test with Dunn's multiple comparisons test.

*p<0.05, **p<0.01

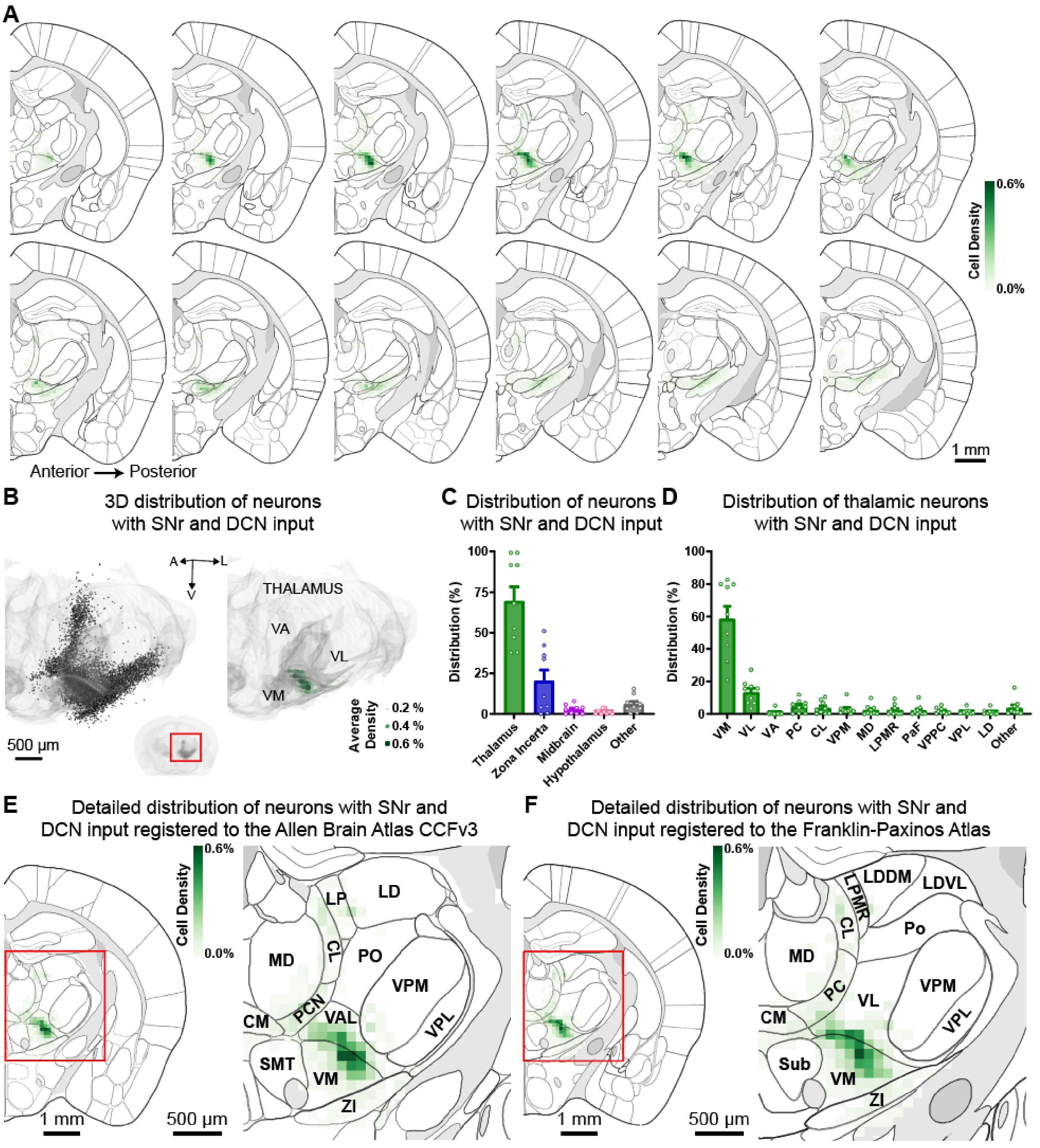


**Roth et al. Figure S2**

**Figure S2 (related to Figure 2): Registration of thalamic neurons with SNr and DCN inputs to the Franklin-Paxinos Atlas.**

(A) Serial 2D coronal sections at 100-μm intervals of the Franklin-Paxinos Atlas. The distribution of labeled neurons with SNr and DCN inputs is shown as the average relative density in 100 × 100 μm bins (green, see methods) from n=9 mice.

(B) 3D projection of the thalamus with individual labeled neurons from all 9 mice overlayed (left) and the average relative density of labeled neurons in 100 × 100 × 100 μm bins (right). Thalamus volume in light gray, VM, VL, and VA highlighted in dark gray. Green circles denote average relative density. Inset shows 3D reconstruction of the entire brain with the enlarged region marked with a red rectangle.

(C) Distribution of neurons with SNr and DCN inputs across brain regions according to the Franklin-Paxinos Atlas. Bars denote mean and circles individual mice. Error bars, SEM. n=12,181 cells from 9 mice.

(D) Distribution of neurons with SNr and DCN inputs within the thalamus according to the Franklin-Paxinos Atlas. Individual thalamic nuclei with an average of ≥1% of labeled cells, as well as VA, are listed, remaining nuclei are combined as "other". Bars denote mean and circles individual mice. Error bars, SEM. n=7,514 cells from 9 mice.

(E) Detailed distribution of neurons with SNr and DCN inputs at a representative 2D coronal section. Enlarged view of area marked with a red rectangle with brain regions outlined according to the Allen Brain Atlas CCFv3.

(F) Detailed distribution of neurons with SNr and DCN inputs at a representative 2D coronal section. Enlarged view of area marked with a red rectangle with brain regions outlined according to the Franklin-Paxinos Atlas.

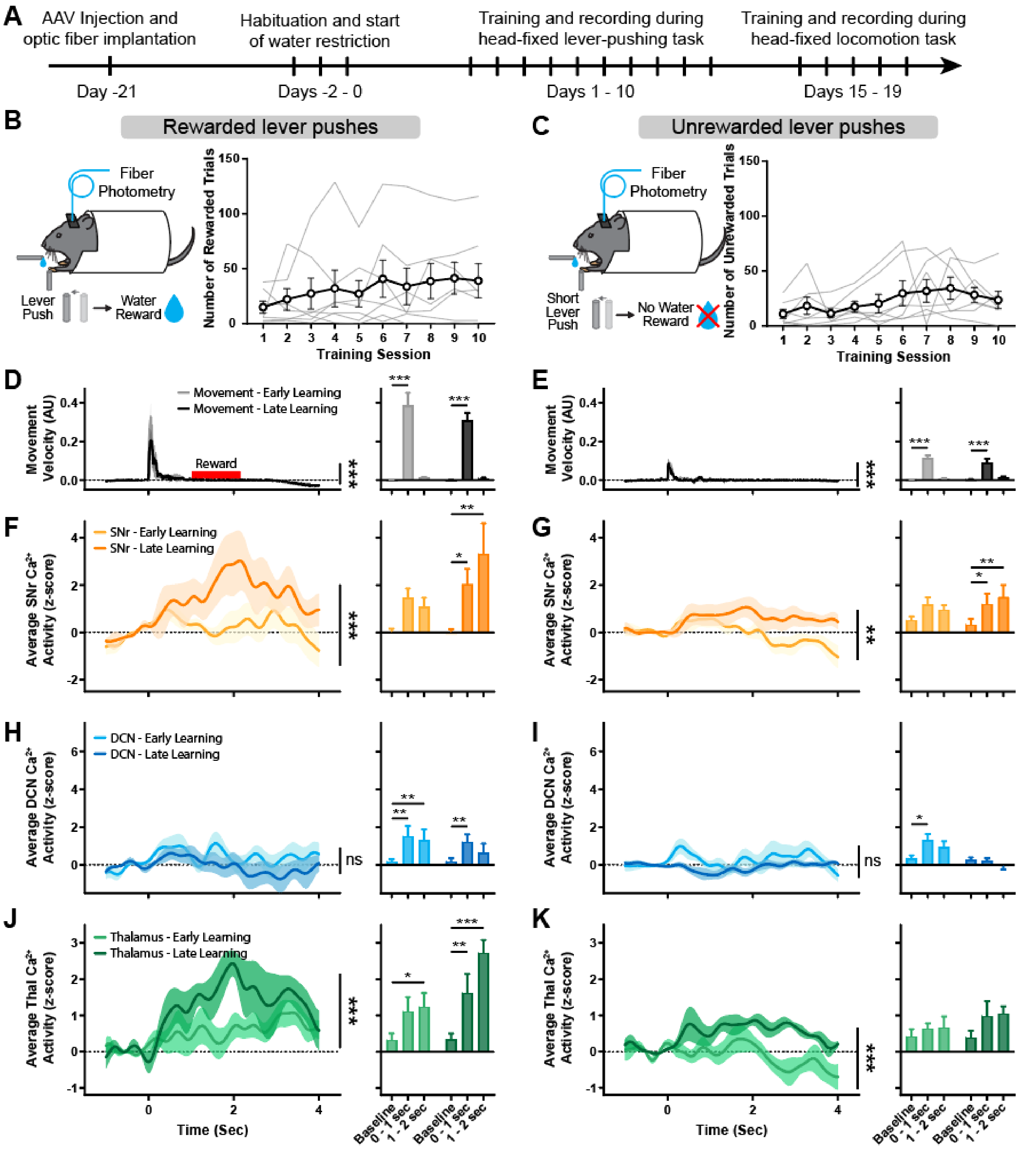


**Roth et al. Figure S3**

**Figure S3 (related to Figure 3): Multi-site $Ca^{2+}$ fiber photometry of SNr, DCN, and thalamus during rewarded and unrewarded trials of lever-pushing task.**

(A) Timeline of surgeries and behavior experiments for fiber photometry recordings of SNr, DCN, and thalamus neurons with dual inputs.
(B) Left: Schematic of mouse performing rewarded lever-push. Right: Number of rewarded lever pushes in mice learning the lever-pushing task over the course of 10 days. Thin grey lines represent individual mice and bold line denotes mean. Error bars, SEM.
(C) Left: Schematic of mouse performing unrewarded lever-push. Right: Number of unrewarded lever pushes in mice learning the lever-pushing task over the course of 10 days. Thin grey lines represent individual mice and bold line denotes mean. Error bars, SEM.
(D) Left: Average velocity of lever movements aligned to movement onset of rewarded trials during early (grey) and late (black) training sessions. Solid lines represent mean and shaded areas denote SEM. n=7 mice. p<0.001, two-way ANOVA interaction. Right: Average maximal movement velocity at baseline, during movement (0-1 sec), and after movement (1-2 sec). Bars denote mean. Error bars, SEM. Two-way ANOVA with Holm-Sidak multiple comparisons test.
(E) Left: Average velocity of lever movements aligned to movement onset of unrewarded trials during early (grey) and late (black) training sessions. Solid lines represent mean and shaded areas denote SEM. n=7 mice. p<0.001, two-way ANOVA interaction. Right: Average maximal movement velocity at baseline, during movement (0-1 sec), and after movement (1-2 sec). Bars denote mean. Error bars, SEM. Two-way ANOVA with Holm-Sidak multiple comparisons test.
(F) Left: Average z-scored $Ca^{2+}$ activity in SNr aligned to lever movement onset of rewarded trials during early (light orange) and late (dark orange) training sessions. Solid lines represent mean and shaded areas denote SEM. n=7 mice. p<0.001, two-way ANOVA interaction. Right: Average maximal z-scored $Ca^{2+}$ activity at baseline, during movement (0-1 sec), and after movement (1-2 sec). Bars denote mean. Error bars, SEM. Two-way ANOVA with Holm-Sidak multiple comparisons test.
(G) Left: Average z-scored $Ca^{2+}$ activity in SNr aligned to lever movement onset of unrewarded trials during early (light orange) and late (dark orange) training sessions. Solid lines represent mean and shaded areas denote SEM. n=7 mice. p=0.003, two-way ANOVA interaction. Right: Average maximal z-scored $Ca^{2+}$ activity at baseline, during movement (0-1 sec), and after movement (1-2 sec). Bars denote mean. Error bars, SEM. Two-way ANOVA with Holm-Sidak multiple comparisons test.
(H) Left: Average z-scored $Ca^{2+}$ activity in DCN aligned to lever movement onset of rewarded trials during early (light blue) and late (dark blue) training sessions. Solid lines represent mean and shaded areas denote SEM. n=7 mice. p>0.999, two-way ANOVA interaction. Right: Average maximal z-scored $Ca^{2+}$ activity at baseline, during movement (0-1 sec), and after movement (1-2 sec). Bars denote mean. Error bars, SEM. Two-way ANOVA with Holm-Sidak multiple comparisons test.
(I) Left: Average z-scored $Ca^{2+}$ activity in DCN aligned to lever movement onset of unrewarded trials during early (light blue) and late (dark blue) training sessions. Solid lines represent mean and shaded areas denote SEM. n=7 mice. p=0.632, two-way ANOVA interaction. Right: Average maximal z-scored $Ca^{2+}$ activity at baseline, during movement (0-1 sec), and after movement (1-2 sec). Bars denote mean. Error bars, SEM. Two-way ANOVA with Holm-Sidak multiple comparisons test.
(J) Left: Average z-scored $Ca^{2+}$ activity in thalamus neurons with SNr and DCN inputs aligned to lever movement onset of rewarded trials during early (light green) and late (dark green) training sessions. Solid lines represent mean and shaded areas denote SEM. n=7 mice. p<0.001, two-way ANOVA interaction. Right: Average maximal z-scored $Ca^{2+}$ activity at baseline, during movement (0-1 sec), and after movement (1-2 sec). Bars denote mean. Error bars, SEM. Two-way ANOVA with Holm-Sidak multiple comparisons test.
(K) Left: Average z-scored $Ca^{2+}$ activity in thalamus neurons with SNr and DCN inputs aligned to lever movement onset of unrewarded trials during early (light green) and late (dark green) training sessions. Solid lines represent mean and shaded areas denote SEM. n=7 mice. p<0.001, two-way ANOVA interaction. Right: Average maximal z-scored $Ca^{2+}$ activity at baseline, during movement (0-1 sec), and after movement (1-2 sec). Bars denote mean. Error bars, SEM. Two-way ANOVA with Holm-Sidak multiple comparisons test.

*p<0.05, **p<0.01, ***p<0.001

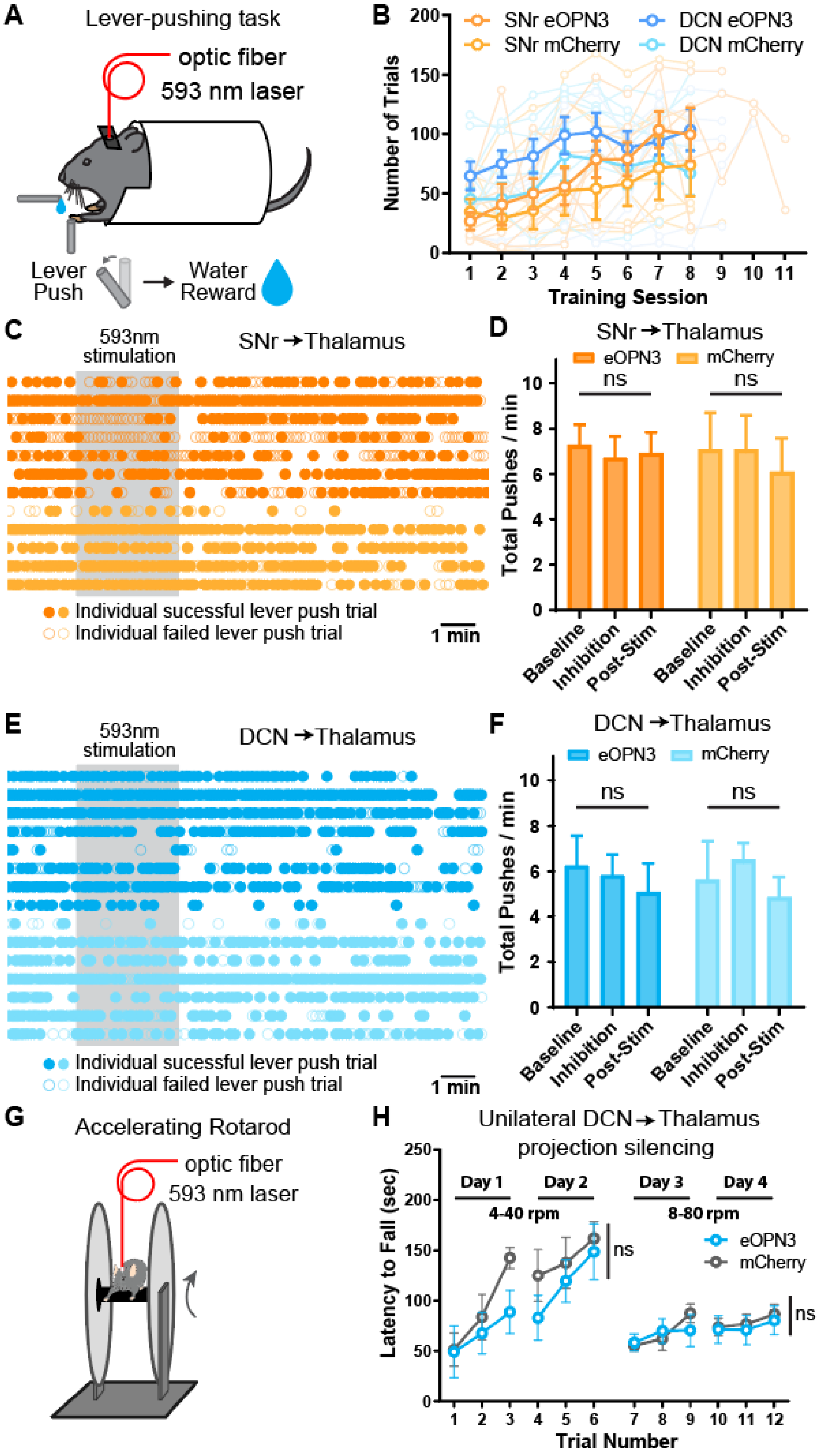


**Roth et al. Figure S4**

**Figure S4 (related to Figure 4): Optogenetic silencing of SNr to thalamus and DCN to thalamus projections.**

(A) Schematic of mouse performing head-fixed lever-pushing task with optic fiber for optogenetic silencing.
(B) Behavior performance of mice learning the lever-pushing task.
(C) Individual lever-push trials during optogenetic silencing session of mice expressing eOPN3 in SNr (dark orange) and control mice expressing mCherry (light orange). Inhibition period shown with gray shading. Filled circles denote successful lever-pushes and open circles denote failed lever-pushes. Each row represents one mouse.
(D) Number of total lever pushes of mice expressing eOPN3 in SNr (dark orange) and control mice expressing mCherry (light orange) before (baseline), during (inhibition), and after (post-stim) optogenetic silencing of SNr to thalamus projections. Silencing was performed in mice well-trained on the head-fixed lever-pushing task. Bars denote mean. Error bars, SEM. eOPN3: n=7 mice; mCherry: n=5 mice. Two-way ANOVA with Holm-Sidak multiple comparisons test.
(E) Individual lever-push trials during optogenetic silencing session of mice expressing eOPN3 in DCN (dark blue) and control mice expressing mCherry (light blue). Inhibition period shown with gray shading. Filled circles denote successful lever-pushes and open circles denote failed lever-pushes. Each row represents one mouse.
(F) Number of total lever pushes of mice expressing eOPN3 in DCN (dark blue) and control mice expressing mCherry (light blue) before (baseline), during (inhibition), and after (post-stim) optogenetic silencing of DCN to thalamus projections. Silencing was performed in mice well-trained on the head-fixed lever-pushing task. Bars denote mean. Error bars, SEM. eOPN3: n=8 mice; mCherry: n=7 mice. Two-way ANOVA with Holm-Sidak multiple comparisons test.
(G) Schematic of mouse performing accelerating rotarod task with optic fiber for optogenetic silencing.
(H) Performance on an accelerating rotarod task of mice expressing eOPN3 in DCN unilaterally (dark blue) and mice expressing mCherry (light blue). DCN to thalamus projections were unilaterally silenced during each training session. Circles denote mean. Error bars, SEM. eOPN3: n=6 mice; mCherry: n=4 mice. Two-way ANOVA, separately for days 1-2 and days 3-4.

ns: not significant
